# MULTIPLE DEFECTS IN MOUSE STEM CELL BASED EMBRYO MODELS LACKING ALL HOX FUNCTION

**DOI:** 10.64898/2026.06.19.733340

**Authors:** Hocine Rekaik, Lucille Lopez-Delisle, Alexandre Mayran, Yuliia Romaniuk, Cynthia Diabanza, Anne-Catherine Cossy, Julie Gauvain, Denis Duboule

## Abstract

Mammals have four genomic copies of an ancestral Hox cluster, with a total of 39 genes. While gene inactivation approaches and full cluster deletions have revealed their critical functions during early development, the effect of removing all Hox function has remained elusive due to both biological and technological challenges. We have used mouse gastruloids, an ES cells-derived embryo model where Hox genes are properly activated in time and space, to assess the effect of their complete absence. We report that gastruloids lacking Hox function can still elongate and reach a general shape resembling their control counterparts, with a well-established AP polarity. However, unlike controls, they fail to produce any endoderm and are unable to properly segment their presomitic mesoderm into persistent somite-like structures. Instead, they produce a type of mesoderm with a more anterior identity. We thus propose that, in this system at least, HOX proteins are necessary to posteriorize an existing anterior ‘ground-state’ structure, in part by promoting and/or maintaining the epithelialization of cellular condensations. Multiomes analysis revealed range of modifications in chromatin accessibility in the absence of any HOX proteins, involving in particular variations in the binding of the co-factor PBX1. In contrast, neuro-mesodermal progenitor (NMP) cells are not overtly affected in mutant gastruloids, even though they normally initiate strong Hox gene transcription, suggesting that these cells are used as vehicles to translate a temporal sequence of activation into an AP colinear transcript distribution, which becomes functional at a later stage only.

## INTRODUCTION

*Hox* genes are a subset of those few hundred vertebrate genes containing a homeobox (see refs in^1^). This DNA sequence encodes a peptide that can bind DNA through an alpha helix that recognize a rather degenerate motif, often in combination with co-factors^2^. In mammals, 39 *Hox* genes are found at four distinct genomic loci, tightly clustered, as a result of the two rounds of genome duplication experienced early on at the roots of the vertebrate lineage^3^. Their initial genetic characterization in *Drosophila* revealed that their mutation induces a morphological transformation of one structure into the likeliness of another one^4,5^, a phenomenon referred to as *homeosis*^6^ and hence the term ‘homeotic genes’ from which ‘*Hox’* and ‘homeobox’ were derived. In flies, these genes were also called ‘selectors’^7^ due to their capacity to select between different fates, for example within imaginal disks^8^.

Functional inactivation of vertebrate *Hox* genes generally complied with the concept of homeosis, yet not entirely. Early on, indeed, phenotypes were interpreted as homeotic transformations, in particular when vertebrae or branchial derivatives were considered^9,10^. Subsequently, however, single or combined mutations suggested that an entire structure -or parts thereof- was lost, rather than transformed^11–14^. Also, two main conceptual frameworks were proposed to account for the mode of action of these proteins, either through a combinatorial system^15^, or through the functional prevalence of the posterior-most expressed gene^16^, often raising difficulties in the interpretation of null phenotypes.

In *Drosophila*, the presence of a single complement of *Hox* gene allowed for a full - though localized- functional depletion, which revealed the existence of a ‘ground state’ segmental structure including elements from head and thoracic segments^4,17^. Likewise, a short and non-segmented fly leg is produced in the absence of all *Hox* function, which may illustrate the ground state used by homeotic proteins to elaborate various ventral appendages^18^. In vertebrates, while the existence of a ground state structure was observed in branchial arches colonized by *Hox*-depleted neural crest cells^19,20^, the deletion of entire *Hox* clusters in isolation mostly strengthened gene-specific phenotypes^21–24^. Combined deletions of several clusters started to reveal as yet invisible phenotypes^20,25^, yet the reproductive problems induced in these mutants hampered the production of animals lacking more than two entire clusters, even at embryonic stages. As a result, the effect of removing all *Hox* functions from an amniote embryo remains unknown.

Recent developments in the production of ES-cells derived embryo models now allows to re-investigate this question. Amongst such models are gastruloids^26,27^, which are elongated structures related to the posterior part of the developing embryo^28^, i.e., the place where *Hox* genes are transcriptionally activated^29^ and where they specify segmental identities in coordination with the segmentation clock^30^. Because gastruloids are entirely derived from ES cell aggregates, high amounts of mutant specimens can be readily obtained (e.g.^31^). We thus set up to delete all *Hox* clusters in a homozygous configuration and to use such *Hox*-less ES cells to produce gastruloids, using as readouts a variety of physical, cellular and molecular parameters. Here we report that *Hox*-less gastruloids can elongate and thus globally resemble in shape control specimen. However, they do not produce any endodermal tissue and fail to segment into mature, somite-like structures. In addition, the mesoderm produced seems to be of an anterior type, suggesting that HOX proteins are used to successively posteriorize the somites and their derivatives. By comparing control and mutant mesoderm cellular trajectories, we conclude that *Hox* gene function is implemented in somitic mesoderm, i.e., much after their initial transcriptional activation in NMP cells.

## RESULTS

We generated a mouse embryonic stem cell line that carries the combined homozygous deletions of the four *Hox* gene clusters (*HoxA, B, C* and *D*; Figure 1a). This *HoxB^−/−^:HoxA^−/−^:HoxD^−/−^:HoxC^−/−^*cell line (referred to below as *Hox^−/−^*) was used to generate gastruloids following a well-established protocol^28^, which triggers a primitive streak-like process and an elongating structure resembling the tail bud (Figure 1b). At 120h after aggregation (120h), control elongating gastruloids present an anterior-posterior organization visible through a patterned expression of *Hox* transcripts similar to the embryo and the presence of axial progenitors (neuro-mesodermal cells or NMPs), which differentiate to fuel both somitic and neural tube tissue (Figure 1c). Gastruloids also produce endodermal, gut-like tissue, though this differentiation pathway is generally less robust.

**Figure 1.**
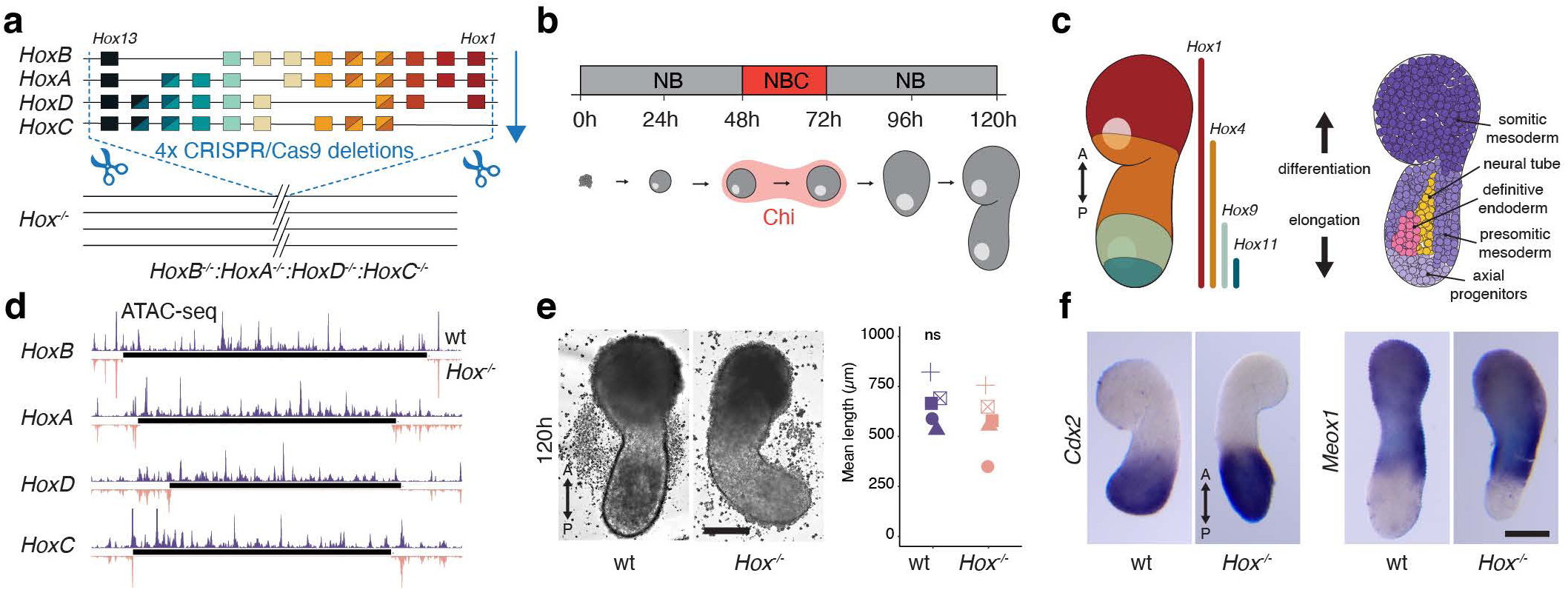
ES-cell-derived gastruloids lacking all HOX function. **a;** Scheme of the four mouse *Hox* clusters showing their sequential deletions in mES cells, leading to full *Hox* mutant (*Hox^−/−^*) cells. Colors illustrate groups of paralogy. **b;** Gastruloid cultures starting from mES cells aggregates up to fully elongated form at 120h. Chiron (Chi) treatment is given between 48h and 72h. **c;** Schematic view of colinear *Hox* genes expression along the anterior (A) to posterior (P) axis in control (wt) gastruloids at 120h (left), with a representation of a rather generic tissue organization at this stage (right). **d;** ATAC-seq profiles covering the four *Hox* clusters in control (purple, above) and *Hox^−/−^* (orange, below) ES cells. Black lines represent the *Hox* clusters mm10 coordinates (*HoxA*: chr6:52155367-52260880; *HoxB*: chr11:96194361-96368254; *HoxC*: chr15: 102921131-103036856 and *HoxD*: chr2:74668310-74765142). The deletions of DNA intervals including the various *Hox* cluster is confirmed by the absence of signals (below). In both *HoxA* and *HoxD* clusters, ATAC signals are detected over parts of the *Hoxa1*, *Hoxa13* and *Hoxd13* genes, which are functionally inactivated (see Extended Data Figure 1b and Methods). **e;** Representative brightfield images of elongated control and *Hox^−/−^*gastruloids at 120h (left), with a quantification of the average lengths showing no significant variation between the two genotypes (right). 5 independent batches were quantified (total gastruloids n = 251 wt, *versus* 379 *Hox^−/−^*). Plotted data represent the mean for each batch and genotype. p-value is determined by unpaired two-sided t-test. **f;** WISH showing the expression of *Cdx2* (left) and *Meox1* (right) in control and *Hox^−/−^* gastruloids at 120h, illustrating the conserved anterior-posterior polarity in mutant gastruloids. Scale bar: 200 µm.

### *Hox^−/−^* mutant gastruloids

Mutant ES cells were morphologically undistinguishable from their wt counterparts (Extended Data Fig. 1a) and the occurrence and precision of all deletions were confirmed either by Sanger sequencing (Extended Data Fig. 1b), or by mining an ATAC-seq dataset obtained from *Hox^−/−^* ES cells (Figure 1d; Extended Data Fig. 1c). We also compared RNA-seq datasets of mutant *versus* control ES cells to make sure that no major modifications had occurred in transcriptomes, since *Hox* genes are either not expressed in ES cells, or at low levels, for some of them. As expected, the transcriptome of *Hox^−/−^* ES cells was comparable to control (Extended Data Fig. 1d, e), supporting the absence of any major *Hox* function before the onset of gastrulation. We next generated *Hox^−/−^* mutant gastruloids, which appeared elongated, with an overall shape globally similar to controls, if anything slightly reduced in their elongation length (ca. 13%, Figure 1e). Likewise, whole mount *in situ* hybridization (WISH) revealed comparable expression domain for the *Cdx2* and *Meox1* genes in *Hox^−/−^* mutants, illustrating a correct distribution of the posterior and anterior regions, respectively (Figure 1f). We thus concluded that *Hox* gene function is not critical for the elongation process, at least in gastruloids.

We compared the differentiation processes and its dynamics between *Hox^−/−^* and control gastruloids by generating single-cell RNA-seq (scRNA-seq) datasets at 72h, 96h, 120h and 144h (Figure 2a). The projected UMAP colored either by timepoints, or by genotypes (Figure 2b, c), already showed variations in cellular signatures at 72h, pointing to an effect of *Hox* genes ablation, even though only a small number of anterior *Hox* genes are normally expressed at this early stage (Extended Data Fig. 2a, left column). In 96h, 120h and 144h gastruloids, similar differences were observed between controls and *Hox^−/−^* mutant specimens, indicating that despite an apparently maintained elongation process, the identity of those cells involved was not identical to controls. We characterized these differences by annotating the various cell clusters using known gene markers either of tissues present in control gastruloid, or extracted from the mouse embryo scRNA-seq atlas^32^, and thus identified clusters enriched in *Hox^−/−^*gastruloids (Figure 2d-f).

**Figure 2.**
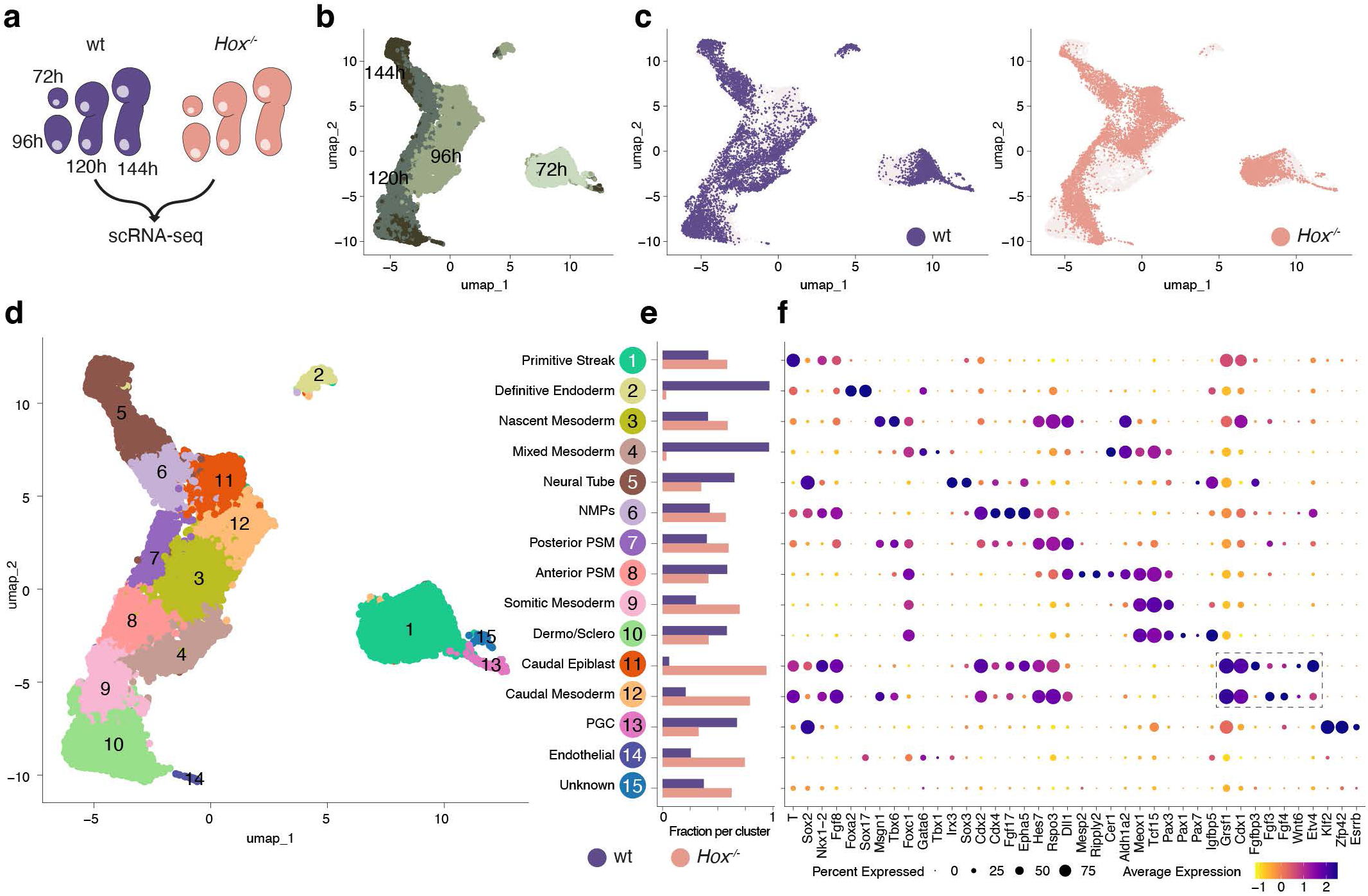
scRNA-seq-based distribution of cell types in *Hox* mutant gastruloids over time. **a;** The various scRNA-seq dataset produced from control (wt) and *Hox^−/−^* mutant gastruloids at 72h (n=2), 96h (n=2), 120h (n=2) and 144h (n=1). **b**; UMAP of scRNA-seq data with control and *Hox^−/−^* mutants cells colored according to their developmental stage. **c;** UMAP of control (purple, left) and *Hox^−/−^* mutants (orange, right) cells showing a largely complementary distribution of cells for the two genotypes. **d;** UMAP showing cells colored by unsupervised clusters. The tentative cellular identity for each cluster is shown on the right. **e;** Fraction (%) of control and *Hox^−/−^* mutant cells in each cellular cluster described under **d**. **f;** Dotplot of selected genes enriched in each cluster dispatched under **d**. The size of the dots represents the proportion of cells expressing any given gene (indicated below) in each cluster whereas colors represent levels of expression (low, yellow; high, blue).

At 72h, the vast majority of cells of both genotypes had a primitive streak-like identity. Indeed, most genes used as markers for primitive streak were not transcriptionally affected in mutant specimen. Few genes only showed a significant difference in expression between the two conditions (Extended Data Fig. 2b), which likely explains the shift observed in the UMAP. At that early stage, the absence of any *Hox* gene function thus induced a rather limited change in gene expression altogether. In 96h gastruloids, however, the shift in cell identities was clear, with several cellular populations affected in mutants. For instance, ‘mixed mesoderm’ (including cardiopharyngeal and cranial mesoderm) was absent (Figure 2c-e, cluster 4), as well as definitive endoderm, a tissue often -yet not always- produced in control gastruloids (Figure 2c-e, cluster 2), suggesting a direct role for *Hox* genes in its differentiation.

### Impairment of endoderm and notochord differentiation

We further assessed the absence of endoderm markers (e.g., *Sox17, Foxa2*) in our scRNA-seq and confirmed this deficiency in endoderm formation by RT-qPCR in several batches of mutant gastruloids, while such transcripts were present in 4 out of 6 control batches (Extended Data Fig. 3a-d). We also observed this absence of endoderm by using an *in vitro* assay to differentiate ES cells towards definitive endoderm by using combined activin A and *Wnt* treatments^33,34^. Under these conditions, unlike control ES cells, *Hox^−/−^* ESC showed only low levels of *Foxa2* or *Sox17* transcripts (Extended Data Fig. 3b and 3c). To substantiate these results, we produced gastruloids treated with Activin A, using a protocol whereby more anterior tissues are produced, with an enrichment in endodermal contributions^35^ (Figure 3a).

**Figure 3.**
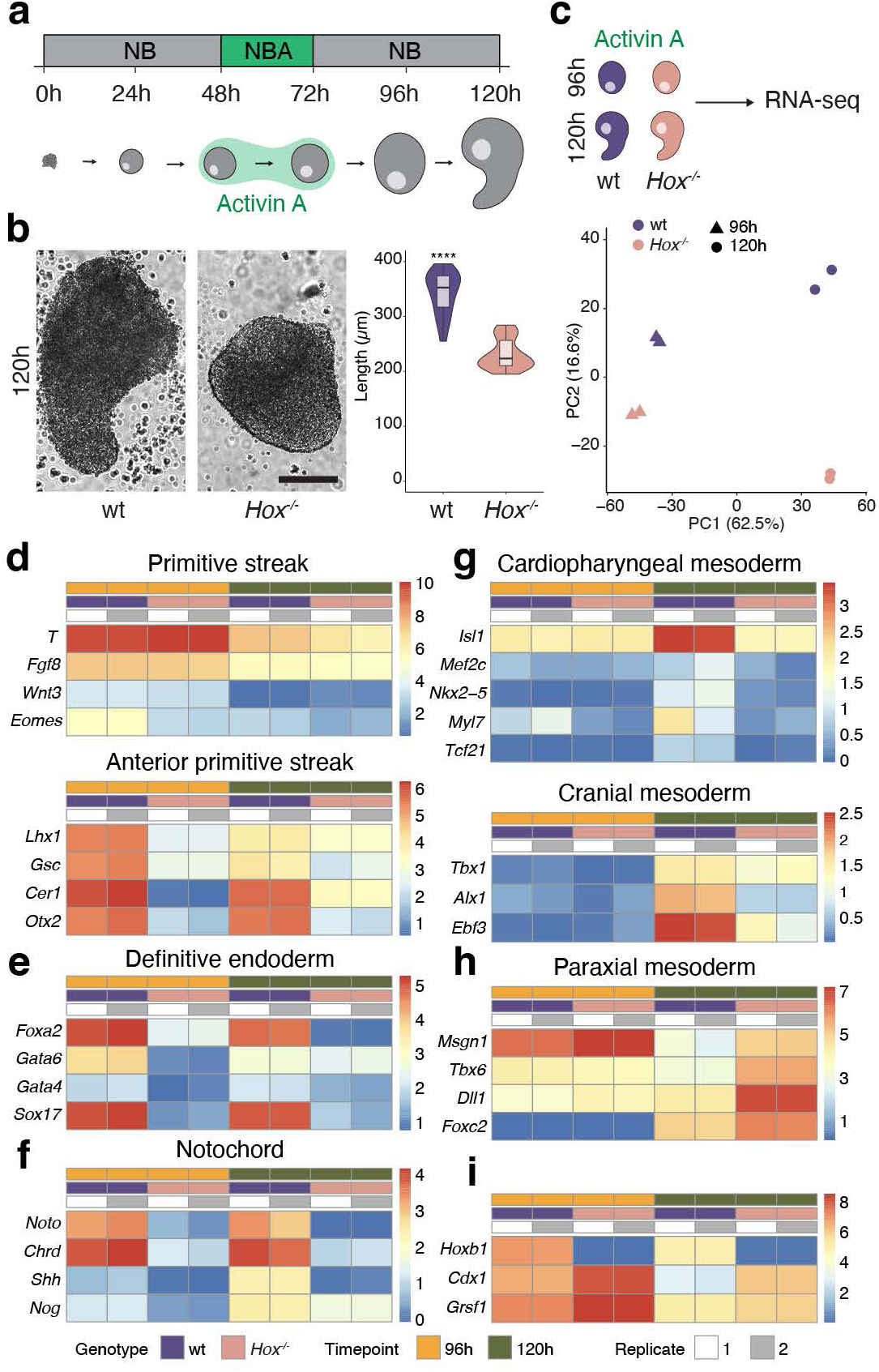
*Hox* gene function is required for anterior patterning and growth in Activin A-induced organoids. **a;** Schematic of the Activin A-based differentiation protocol used to generate anteriorized gastruloids from mES cell aggregates. **b;** Representative images (left) and quantification (middle) showing that *Hox^−/−^* mutant organoids are significantly shorter and less elongated than wt controls at 120h. n = 30. p-value is determined by unpaired two-sided t-test (****P < 0.0001), a defect not seen in Chiron-based gastruloids. Box plots with median value and 25–75% percentiles, whiskers represent minimum and maximum values no further than 1.5 * IQR from the hinge. **c;** Experimental design for bulk RNA-seq analysis of wt and *Hox^−/−^* Activin-treated gastruloids at 96h and 120h (top). Principal component analysis (PCA) (bottom), with a clear transcriptional separation by genotype on the second component. **d–i;** Heatmaps of selected RNA-seq data reveal distinct tissue-specific gene expression profiles. *Hox^−/−^* gastruloids fail to robustly activate markers of definitive endoderm, cardiopharyngeal mesoderm and notochord. Conversely, they show an upregulation of markers associated with paraxial mesoderm and specific transcripts such as *Cdx1* and *Grsf1*.

Comparative bulk RNA-seq of wt gastruloids showed a significant increase of anterior markers expression in Activin *versus* Chiron-treated conditions, including markers for definitive endoderm, notochord, cardiopharyngeal and cranial mesoderm (Extended Data Fig. 4a). The anterior identity of Activin gastruloids was confirmed with the preferred expression of anterior *Hox* genes (group 1-2) and a significant delay in the activation of posterior genes, even at 120h (Extended Data Fig. 4b). *Hox* mutant Activin A gastruloids were significantly smaller than wt controls (Figure 3b), an effect not seen after Chiron treatment. Bulk RNA-seq analyses of such mutant Activin gastruloids at 96h and 120h confirmed major differences in RNAs content when compared to wt controls (Figure 3c-i; Extended Data Fig. 4c, d), in particular with a strong reduction of transcripts markers for anterior primitive streak, definitive endoderm and notochord. Likewise, markers of cardiopharyngeal mesoderm tissue (part of cluster 4 on Figure 2c-e) were much weaker at 120h, confirming the result observed in Chiron treated gastruloids^36,37^.

Of note, both the cardiopharyngeal mesoderm and the endoderm share the same developmental window and spatial origin in the anterior streak of developing embryos^38,39^, which is distinct from the posterior primitive streak, as shown by markers such as *T, Gsc* and *Eomes*. The quasi-absence of anterior primitive streak markers in *Hox* mutant Activin gastruloids (Figure 3d) suggested that an impairment in patterning of the primitive streak tissue in such mutant specimens may hamper the differentiation of both cardiopharyngeal mesoderm and endoderm. In addition to supporting a lack of endoderm in *Hox* mutant gastruloids, these Activin treated specimen were also unable to produce notochord cells, at least to the amount normally detected under these conditions (Figure 3f).

### Defective somite formation

To assess whether mesodermal derivatives other than cardiopharyngeal mesoderm and notochord were affected by the lack of *Hox* function, we cultured our gastruloids in Matrigel, which triggers somite formation^40^ (Figure 4a). Under these conditions, a clear morphological transition occurs between the presomitic mesoderm (PSM) and a segmented tissue that contains bulges resembling genuine epithelialized somites (Figure 4b). This segmentation process is comparable in many respects to the output of the segmentation clock, the transition occurring at the determination front^40^, partially corresponding to the rostral ‘end’ of each oscillation^41^. In control gastruloids, newly formed somites display a distinct polarization, as well as signs of anterior-posterior patterning. In contrast, *Hox^−/−^*gastruloids failed to show this type of morphology (Figure 4c, black arrowheads in wt). In contrast, they formed significantly more ectopic outgrowths at the posterior pole where NMPs are located (Figure 4c, white arrowheads, Movie 1; for quantification, see method section). Phalloidin staining confirmed the absence of somite epithelization in the anterior part of mutant gastruloids, whereas some pseudo-polarized somites, often ill-formed, were observed right above the PSM-SM transition, suggesting that the epithelialization process was initiated, but either not completed or not maintained (Figure 4d and e). As a consequence, the clear and regular epithelialized rosettes observed in wt specimen, corresponding to strong foci of actin staining, were rare and/or ill-formed in the main part of the mutant counterparts (Figure 4f).

**Figure 4.**
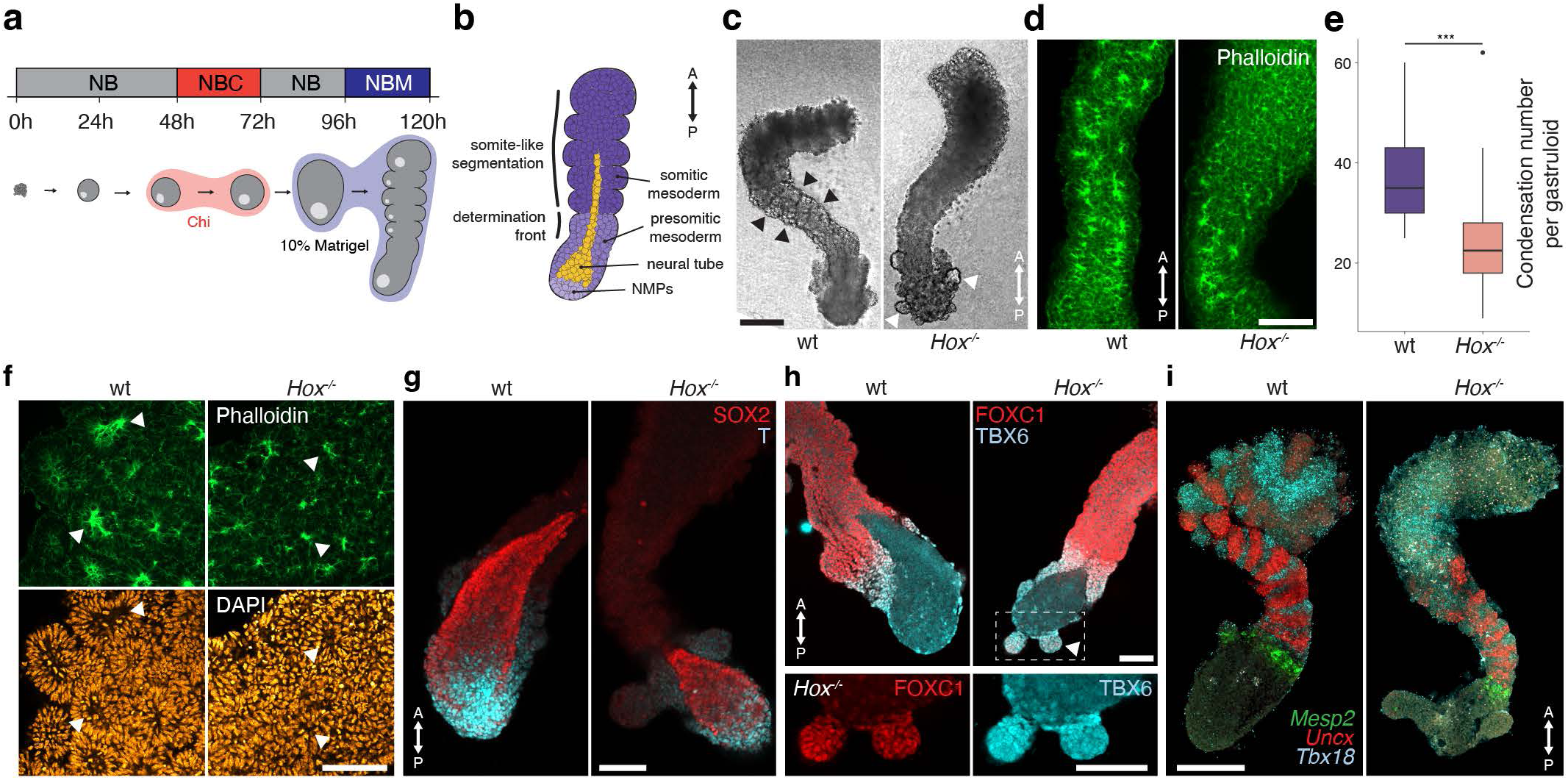
Defective segmentation in *Hox*-less gastruloids. **a;** Experimental scheme of gastruloids transferred onto 10% Matrigel between 96h and 120h. NB, gastruloid medium; C, Chiron; M, matrigel. **B;** Schematic of a 120h gastruloid grown under these conditions, with a generic tissue distribution. The epithelialization of somites occurs first at the level of the transition between PSM and SM. **c;** Comparison between control and *Hox^−/−^*gastruloids exposed to Matrigel, at 120h. Epithelialized somites are clearly detected in controls (black arrows), while mostly absent from *Hox^−/−^* mutants gastruloids. Ectopic outgrowths are observed at the most posterior aspect of the mutant counterparts (white arrowheads). Scale bar: 200 µm. **d;** Phalloidin immunostaining of actin filaments in 120h gastruloids. Somitic-like condensations are visible in controls throughout the entire structure anterior to the PSM to SM transition (t), whereas only few polarized structures are scored in mutants gastruloids. **e,** Quantification of the number of epithelized structures detected per gastruloid, using phalloidin staining. A mix of two independent quantifications is shown. Box plots with median value and 25–75% percentiles, whiskers represent minimum and maximum values no further than 1.5 * IQR from the hinge. Data beyond the whiskers are plotted individually. n = 27. p-value is determined by unpaired two-sided t-test (***P < 0. 001). **f;** Immunofluorescence of Phalloidin (top) and DAPI (bottom) in 120h control and *Hox^−/−^* gastruloids. The strong reduction of actin-based condensations in mutant gastruloids is highlighted (arrowheads). **g;** Double SOX2 and T immunostainings of 120h control and *Hox^−/−^* gastruloids. NMPs cells (positive for both markers) were detected in both genotypes. **h;** Double TBX6 and FOXC1 immunostainings in 120h gastruloids. TBX6 (PSM) and FOXC1 (somitic cells) signals were observed in control and *Hox^−/−^*gastruloids (top). The ectopic outgrows over-represented in mutant gastruloids were positive for both TBX6 and FOXC1 (bottom). **i;** RNA FISH for *Mesp2*, *Uncx* and *Tbx18* in control and *Hox^−/−^* gastruloids at 120h. Expression of *Mesp2*, corresponding to the PSM to SM transition, was observed in both control and mutant gastruloids. The patterned distribution of *Uncx* and *Tbx18* was not maintained anteriorly in mutant gastruloids, coincident with an absence of epithelialization.

*Hox* mutant gastruloids passed onto Matrigel showed the expected presence of SOX2-and T-positive NMP cells in their posterior most aspects (Figure 4g), as well as cells differentiating towards a PSM identity with FOXC1 and TBX6 proteins, similarly to controls (Figure 4h, top). Of note, the increased ectopic structures observed in mutant specimens expressed these same PSM markers, suggesting an erratic differentiation and/or cellular behaviour from the *Hox^−/−^* NMPs (Figure 4h, bottom). Fluorescence *in situ* hybridization (FISH) for *Mesp2* mRNAs, normally enriched at the S-1 level in PSM, i.e., just before morphological segmentation occurs^42,43^, was identical to control (Figure 4i). However, the *Tbx18* and *Uncx* genes, which are expressed in the anterior and posterior part of the formed somite, respectively, displayed an altered pattern in the mutant, despite the absence of condensations, indicating that the segmentation clock was properly implemented in mutant specimens. While expression resembled the control situation at the PSM to SM transition, patterning was rapidly disrupted in more anterior region, matching the detected loss of epithelization and the presumptive defect in somite differentiation (Figure 4i).

Segmentation of the PSM is controlled by a molecular oscillator involving the *Notch*, *Fgf* and *Wnt* pathways^44^. We assessed the expression of genes members of these pathways and no significant change was observed in 120h mutant gastruloids, neither for the *Wnt*/*Fgf* (*Wnt3a*, *Fgf8* or *Fgf17*), nor for the *Notch* (*Hes7*, *Dll1*, *Notch1* or *Lfng*) pathways (Extended Data Fig. 5a, Supplementary Table 1). However, the *Cer1* gene^45^, which encodes a secreted inhibitor of *Wnt*, *Bmp* and *Nodal* signaling^46^, was significantly downregulated (Extended Data Fig. 5a, b). During somitogenesis, *Cer1* transcripts are found in the most anterior part of newly formed somites^47,48^. CER1 Immunostaining showed a complete loss of signal in mutant gastruloids, whereas a clear band of secreted material was observed at the PSM to SM transition in control specimen (Extended Data Fig. 5c). Markers of the forming anterior somite such as *Epha4* and *Pdch8* were detected in mutant specimens, suggesting that those cells normally expressing *Cer1* were present (Extended Data Fig. 5b, d). The absence of matured somitic mesoderm was also reflected by the clear decrease in *Pax7* expression, a marker of dermomyotome, while the expression of the sclerotome marker *Pax1* was mostly maintained (Extended Data Fig. 5b). The decrease in *Cxcl12* and *Igfbp5* expression (Extended Data Fig. 5a), two genes expressed in embryonic somitic mesoderm^49,50^, supported this observation.

Noteworthy, a weak but clear and penetrant gain of expression of SNAI1 was scored in mutant gastruloids, anterior to its normal expression domain (Extended Data Fig. 5e, dashed line), which was further quantified (Extended Data Fig. 5f). Such an increased expression of *Snai1* may contribute to the impairment of somitic mesoderm cells to transit towards an epithelial identity and shape, a hypothesis that will call for additional evidence. Altogether, it appeared that the mutant ‘somitic’ mesoderm was different in identity from its wt counterpart.

### Lack of mesoderm posteriorization

Amongst the cell types gained in *Hox^−/−^* Chiron gastruloids, we identified two unusual cellular identities resembling the embryonic caudal epiblast and caudal mesoderm, expressing a unique combination of markers such as *Cdx1*, *Grsf1* and *Etv4*. These cells were only scarcely produced in control gastruloids, while highly enriched in 96h mutant specimens (Figure 2e, dashed rectangle; Extended Data Fig. 6a). RNA velocity analysis showed a transition from caudal epiblast to caudal mesoderm cells, suggesting the differentiation from primitive streak-derived progenitor cells expressing *Tbxt (T)* and *Nkx1-2*, towards a paraxial mesoderm-like tissue expressing *Tbx6* and *Msgn1* (Extended Data Fig. 6b, c). Integration with the mouse embryo atlas^32^ (Extended Data Fig. 7a) tentatively associated these *Hox^−/−^* mutant cells to those located in the caudal part of the normal embryo between E7.75 and E8, around the primitive streak, which after ingression generate the most anterior paraxial mesoderm (Extended Data Fig. 7b-d), defined as ‘caudal lateral epiblast’ and ‘paraxial mesoderm C’ in ref.^32^.

These early progenitors are clearly distinct from the neuro-mesodermal progenitors (NMPs) that appear later in the embryo, at ca. E8.25 and at a more posterior position, which generate both the neural tube and the paraxial mesoderm of the following posterior somites (Extended Data Fig. 7d, e). The expression of several markers including *Cdx1* and *Grsf1* was shared by both the caudal epiblast cluster derived from *Hox^−/−^* gastruloids, and the caudal epiblast of the embryo (Extended Data Fig. 8a), whereas amongst all *Hox* genes, only *Hoxa1*, *Hoxb1* and *Hoxb2* transcripts, as well as *Hoxb8* to some extent, were detected in this embryonic tissue, suggesting that *Hox* genes have a rather limited contribution to the maintenance and differentiation of the early caudal epiblast (Extended Data Fig. 8b), likely due to its very anterior position. In contrast, NMPs appearing subsequently in the embryo show a colinear activation of the entire *Hox* gene clusters (Extended Data Fig. 8b), which indicates a possible role for these genes in the NMP-mediated patterning of axial tissue, as previously proposed (e.g.^51^).

### HOX depletion has little effect on gastruloid NMP cells

The global transcriptomic signature of NMPs in *Hox^−/−^*gastruloids was nevertheless unexpectedly similar to that of control NMPs, the main difference being the unsurprising absence of any *Hox* transcripts (Extended Data Figure 9), indicating that the direct effect of HOX proteins in these cells is moderate at best, even though these are the cells where the temporal colinear expression of *Hox* genes occurs. We thus conclude that, at least in gastruloids, NMP cells are only marginally dependent on *Hox* function, despite their critical role in activating this gene family. This close-to-normal status of *Hox* mutant NMP cells is confirmed by their capacity to generate neural tube, presomitic mesoderm and some somitic mesoderm tissue at 120h, thus complying with the apparently normal elongation observed in mutant specimens. However, *Hox^−/−^* gastruloids showed an absence of dermomyotome, while sclerotome markers were globally maintained, which are the most differentiated cell types along the somitic lineage that wt gastruloids are capable to produce using our protocol (Figure 2e, cluster 10). We thus inferred that *Hox* gene expression is not mandatory for the initial formation and maintenance of NMPs and that they deploy their function at a much later stage, during the subsequent differentiation process leading to somite production, maintenance and maturation.

### The timing of HOX function

To have a more direct access to the molecular effect(s) of removing all HOX proteins on the various abnormal mesoderm phenotypes described above, and to evaluate the exact time when these proteins become functionally active, we produced combined scATAC-seq and snRNA-seq (‘multiome’) datasets for both *Hox^−/−^* mutant and control gastruloids at 96h (Supplementary Figure 1) and 120h (Figure 5a). The effects scored at 96h were confirmed and amplified at 120h. By focusing on the NMP to somitic differentiation trajectory at 120h, the comparative *Hox^−/−^ versus* wt integrated cluster analysis revealed a clear divergence in cellular identity beginning around the determination front, or soon after, at the transition between posterior and anterior PSM (Figure 5b, c; Extended Data Fig. 10). The analysis of RNA velocity revealed the inferred differentiation dynamics (Figure 5d), from a rather unified cluster of both mutant and wt NMP cells (Figure 5c, cluster 1), to the clear split between the mutant and wt clusters of distinct somitic mesoderms (Figure 5c, clusters 4).

**Figure 5.**
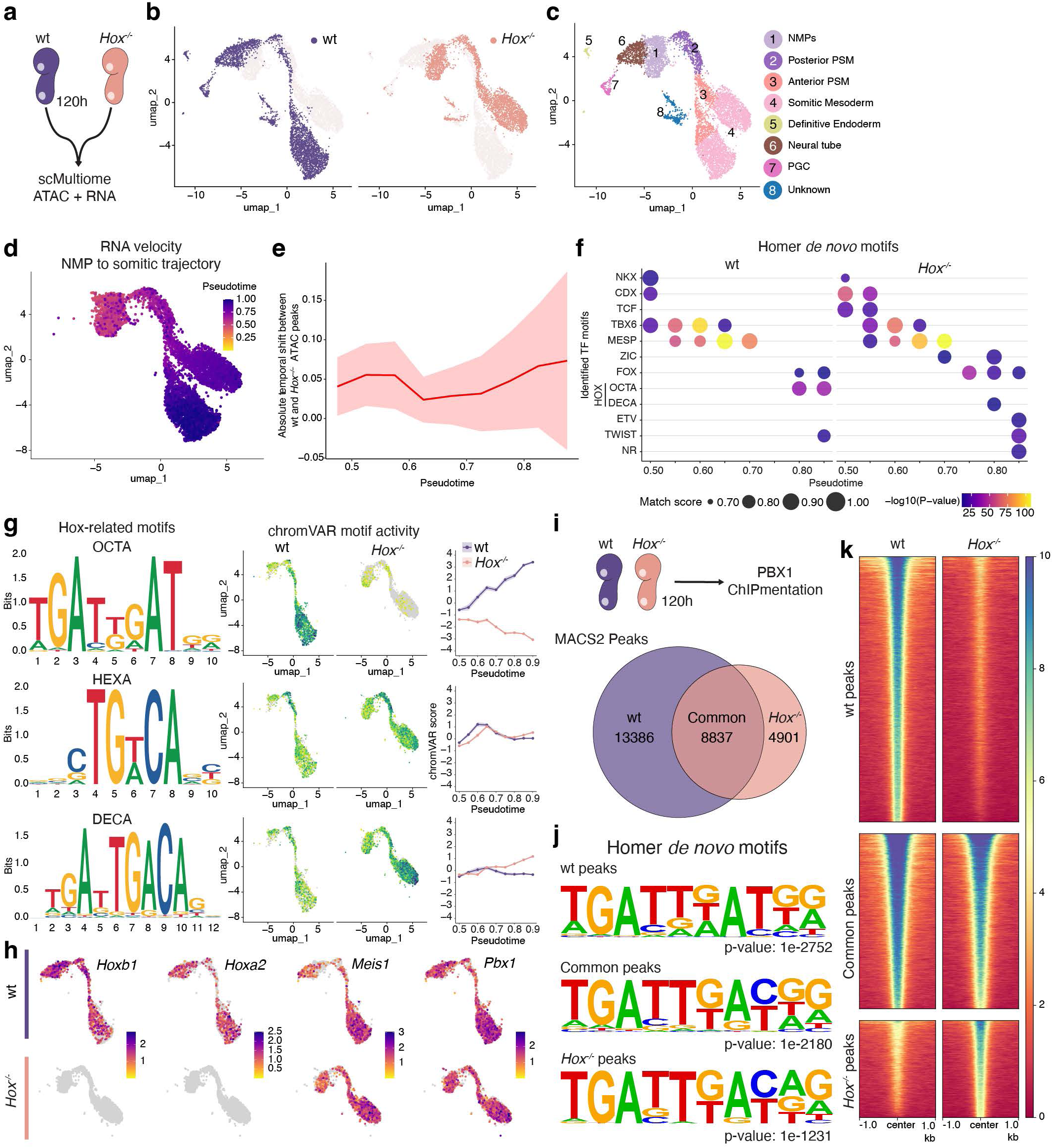
Multiome analysis reveals a redistribution of homeodomain binding sites in *Hox* mutant gastruloids during somite differentiation. **a;** Experimental scheme of 120h control (wt) and *Hox^−/−^* gastruloids used to generate a single-cell multiome dataset, combining both ATAC and RNA analysis. **b;** UMAP projection of the integrated multiome data (RNA and ATAC), with cells colored by genotype (wt, purple; *Hox^−/−^,* orange) showing the distribution of cells at 120h. **c;** UMAP projection showing cells colored by unsupervised clusters, with cluster identities indicated on the right. **d;** UMAP displaying RNA velocity of cells at 120h, revealing the inferred differentiation dynamics. **e;** Plot showing the mean ± SD of the absolute temporal shift between wt and *Hox^−/−^* ATAC peaks over pseudo-groups of cells. Values are measured within the ATAC peak collections representing the most variable peaks in each pseudotime group. Peaks become progressively more divergent in cells that have passed the determination front (i.e., at ca pseudotime 0.7). **f;** Dot plot showing Homer *de novo* motif enrichment for each pseudo-group, with identified transcription factor motifs in both wt and *Hox^−/−^* cells. **g;** On the left, DNA sequences of HOX-related binding motifs (OCTA, HEXA, DECA). Center: UMAPs of chromVAR motif activity for each sequence in the left. Right: quantification of chromVAR motif activity across pseudotime (plot line + ribbon showing mean ± SD). **h;** UMAPs of RNA quantification for *Hox* genes (*Hoxb1*, *Hoxa2*) and their cofactors *Meis1* and *Pbx*, as extracted from the multiome dataset. HOX-related motif activity occurs independently of *Hox* gene and cofactor (*Meis1, Pbx1*) expression. **i;** Scheme showing ChIPmentation of PBX1 in 120h wt and *Hox^−/−^* gastruloids. Below is a Venn diagram showing MACS2 peak distribution. **j;** Homer *de novo* motif logos enrichment either in wt peaks, common peaks or *Hox^−/−^* peaks. **k;** Heatmaps validate the distinct enrichment profiles of these sets of peaks.

Using pseudotime, the differentiation trajectory was divided into series of successive pseudo-groups, starting from NMP cells up to somitic mesoderm (Figure 5d; Extended Data Fig. 10c), and the top enriched ATAC-seq peaks were scored for each pseudo-group, either in wt or in mutant gastruloids. We then plotted the extent of the differences in those ATAC-seq peaks for each pseudo-group points and noted that the difference between the two conditions started to increased significantly at ca. pseudotime point 0.7 (Figure 5e), roughly corresponding to the determination front where somite-like structures start to form. The difference between the two sets of selected ATAC-seq peaks then continued to increase along with the trajectory, reaching a maximum in somitic mesoderm (Figure 5e), indicating that *Hox* gene function became fully functional at this stage, likely to initiate and secure this differentiation pathway.

### Divergence in HOX binding sites accessibility

For each pseudotime group of either wt or mutant cells, we used Homer search for *de novo* motif enrichment within the subsets of ATAC-seq peaks and identified the most significantly represented binding sites (Figure 5f). Along the NMP to PSM trajectory, the overall motif enrichment patterns, such as for TBX6 and MESP, were similar between wt and mutant gastruloids, indicating that the same core developmental program was active. Some exceptions were nevertheless noted. For instance, CDX and TCF motif enrichments were slightly higher in the mutant, which may be explained by the corresponding enrichment of *Cdx1* and *Fgf3* transcripts (Extended Data Fig. 6a). In contrast, the sets of motifs enriched in wt and mutant cells along the PSM to SM were significantly different. This was highlighted, for example, by the specific enrichment of FOX-related motifs in the mutant, suggesting that a distinct developmental program was implemented in this particular developmental trajectory.

For HOX-related motifs, the consensus octamer HOX binding site (OCTA), which is normally bound by HOX proteins dimerized with their TALE co-factor proteins MEIS or PBX (see^2,52^ for references) was enriched in wt ATAC-seq peaks only from the late pseudotime points, thus confirming a functional impact occurring after the PSM to SM transition (Figure 5f, g). In mutant gastruloids, the OCTA site was not amongst the most detected, indicating that, in the absence of HOX proteins, most of their binding sites were no longer accessible, for instance to other homeodomain proteins. In contrast, the enrichment in decamer binding site (DECA), which is normally occupied by the TALE proteins PBX dimerized with their MEIS or PREP partners without HOX participation (e.g.^52^), was slightly enriched in the mutant ATAC-seq peaks, when compared to wt, again at a late pseudotime point (Figure 5f, g). We did not see any variation in the presence of hexamer binding site (HEXA; Figure 5g). These differences occurred without any detectable changes in the transcription of *Meis1* and *Pbx1* (Figure 5h).

These results suggest that, in the absence of any HOX proteins, PBX-containing complexes may be redirected to bind an additional subset of binding sites, including the decamer sequence (mostly defined by the presence of a CA at positions 9-10, figure 5g). To verify this possibility, we looked at the binding profiles of PBX1 in both wt and mutant gastruloids at 120h. More than 13’000 peaks were specific for wt cells, mostly containing OCTA-like binding sites occupied in wt, but no longer accessible in mutant cells (Figure 5i-k). More than 8’000 peaks were found common to both wt and mutant cells and close to 5’000 peaks were specific to the mutant condition. These latter peaks contained a binding sequence related to the decamer consensus, with a large proportion of CA at positions 9-10 (Figure 5i, j). These results suggest that two distinct effects may occur in the absence of HOX proteins: first, a direct impact upon those octamer binding sites normally occupied by HOX/TALE complexes, likely modifying the regulation of several target genes. Secondly, the *de novo* binding of PBX1-containing complexes to a set of binding sites that they would not normally occupy, or occupy with lower probability, which may trigger the mis-regulation of other target genes.

## DISCUSSION

In murine embryos, the removal of all HOX function in some secondary axial structures led to severe truncations, if not a complete agenesis^11,13,14^ and hence the question was raised regarding the persistence of trunk structures in the absence of the entire *Hox* gene complement, i.e., the potential existence of a ground-state structure, as in flies^17,53^. This question has remained elusive due both to technological challenges (deletion of *HoxA* is an early lethal condition^22^) and to an increased infertility of mice lacking such gene clusters^25^. Here, we show that gastruloids lacking HOX function elongate and resemble their control counterparts, morphologically speaking. Considering the comparisons of transcriptomes between temporal series of both gastruloids and murine embryos^28^, we may tentatively extrapolate this result to embryos and conclude that *Hox* gene function may not be essential for at least a partial extension of the trunk axis. Such extrapolations should be considered with caution, however, due to the way gastruloids are produced. Likewise, the absence in mutant specimens of any endoderm, neither in our Activin-treated gastruloids nor in a targeted differentiation protocol *in vitro*, while strongly suggestive of a major impairment, does not formally demonstrate that *Hox*-less embryos would not be able to produce endodermal derivatives.

The production in our mutant gastruloids of a type of mesoderm that resembles a more anterior embryonic mesoderm than that produced in control specimens, suggests that HOX proteins are required to progressively posteriorize the paraxial mesodermal compartment. Of note, the anterior paraxial mesoderm in mammals produces a few pairs of somite-like condensations that are not maintained during subsequent development and which contribute to parts of the basioccipital bone^54^. It is thus tempting to propose that the hereby reported failure to properly segment and maintain somite-like structures in *Hox*-less gastruloids reflects what normally happens in the most anterior paraxial embryonic mesoderm, in a region where only few (if any) anterior *Hox* genes are expressed. This lack of posteriorization of the mesoderm component was also illustrated by the robust gain of expression of *Cdx1* in mutant gastruloids, a gene that is normally expressed in anterior mesoderm during early gastrulation^55^ and whose genetic inactivation induced defects in the cervical and thoracic regions^56^. Therefore, while *Cdx* genes were proposed to positively regulate *Hox* gene expression (e.g.^57^), the *Hox*-dependent posteriorization of the embryonic trunk mesoderm may partly involve the repression of *Cdx1*.

The production of a few genuine segments just rostral to the PSM-SM transition, as judged by both molecular markers and transcriptomic analyses, indicated that the segmentation clock is implemented in the absence of any *Hox* genes. While the size of the mutant segments was somewhat comparable to controls, their total number was difficult to estimate due to their non-persistence. Therefore, a potential involvement of HOX proteins in the termination of the oscillations through repression of *Wnt* signaling^58^ could not be clearly verified, even though *Wnt3a* transcripts were gained in mutant gastruloids at an earlier stage. Altogether, this observation clarifies the long-standing question of the necessary cross-regulation between the segmentation clock and the *Hox* timer^43,59,60^. Indeed, it suggests that, while the *Hox* timer may be controlled by the segmentation clock, HOX proteins have little effect in the implementation of the oscillations, except perhaps for the synchronization of these two temporal processes, an aspect that is difficult to sort out under these experimental conditions. Likewise, the function of HOX13 proteins either to terminate^61,62^ or to trigger^63^ axial extension could not be assessed, for these terminal genes are barely activated under our culture conditions.

In Matrigel-exposed mutant gastruloids, the non-maintenance of well-formed somites is likely associated with their lack of epithelialization. This defect doesn’t seem to derive from the early step of somite formation, for all components necessary for this mechanism seem to be normally functional in mutants, in particular gene members of the *Notch* pathway as well as *Mesp2*^64^. A salient difference observed early on in mutant specimens was the absence of all CER1 protein in the anterior part of the forming somites^47,48^, unlike in the case in control gastruloids. Despite the absence of any segmentation defect in mice lacking *Cer1* function^65,66^, the observed lack of CER1 may reflect a more substantial problem occurring in mutant somitic cells, leading to their subsequent lack of epithelialization. The latter defect may also be due to the ectopic expression of *Snail* in somitic mesoderm, which could trigger these cells to keep their mesodermal morphology at the expense of epithelialization.

Considering previous reports (e.g.^51^), it is somewhat surprising not to see more severe effects in mutant NMP cells, i.e., those cells where the *Hox* timer is implemented (e.g.^67^). Indeed, mutant NMP cells did not appear drastically modified in their gene expression patterns, nor did they generate completely unexpected cell type derivatives. In fact, the clear differences in both gene expression and morphologies occurred later in gastruloid development and one may thus wonder why *Hox* genes are expressed at these early stages, in these particular precursors. The reason is likely to be found in the mechanism necessary to deploy in space the requested AP colinear expression domains, which would make use of the property of NMP cells to progressively fuel the paraxial mesoderm and CNS with their descents while continuing to self-maintain, to properly organize the nested expression domains necessary for segmental identification^68^. In this view, NMPs are used to distribute an information that they themselves do not require and hence HOX proteins may be functionally rather inert there, until the time has come to participate either to the epithelialization of the somitic mesoderm, or to the determination of neuronal subtypes, for example.

Our *a priori* unbiased multiomes approach confirmed the rather late implementation of HOX function, using binding site accessibility as a proxy, and showed that most of these sites becomes accessible only once they are bound by HOX/TALE complexes. It also revealed that the presence of HOX proteins somehow prevents the binding of TALE multimers to the full variety of available binding sites, since *Hox* mutant gastruloids at 120h showed several thousand of sites bound by PBX1, which were not (or at best much weakly-) bound in the wt counterpart. This may indicate that HOX proteins either outcompete PBX1 binding to decamer sites by driving the multimer complexes towards octamer binding sites or, alternatively, that such complexes simply sequester PBX proteins, titrating them from binding to DECA sites. PBX proteins multimerized with other TALE partners have their own functional importance, including in very anterior parts of the developing embryo^69^. It is thus possible that the absence of HOX proteins leads to PBX proteins to maintain an anterior, normally ‘HOX-free’ program throughout the developing gastruloid. The successive addition of HOX proteins along the developing trunk would then progressively re-direct TALE proteins function towards more posterior programs (see ref.^69^). In this context, HOX proteins, rather than their TALE interactors, could be considered as the necessary co-factors (see^70^) to posteriorize the body in a successive manner, along with trunk extension.

## MATERIALS AND METHODS

### Generation of *Hox^−/−^* mES cells

The *Hox^−/−^* cell line was generated from wild-type mES cells (EmbryoMax 129/SVEV) following the CRISPR/Cas9 genome editing protocol as reported in^71^. Sequential targeting of the *HoxB*, *HoxA*, *HoxD* and *HoxC* clusters (in this sequence) was performed with sgRNAs (Supplementary Table 2) cloned into a plasmid expressing the Cas9-T2A-Puromycin and the U6-gRNA scaffold (gift of A. Németh; Addgene plasmid, 101039). Eight consecutive transfections were performed to obtain homozygous deletions of the four clusters. Each time mES cells were transfected with 5μg of both 5’ and 3’ sgRNA plasmids, then dissociated 48h later for puromycin selection (1.5μg ml^−1^). 5 to 6 days later, clone picking was performed and positive mES cell clones were assessed by PCR screening using specific primers surrounding the targeted region (Supplementary Table 3). Mutations were verified for both alleles by Sanger sequencing (Supplementary Table 4). For the second deletion of the *HoxA* and *HoxD* cluster, sgRNA inside *Hoxa1*, *Hoxa13* and *Hoxd13* were used. The remaining parts of these genes were detected in RNA-seq, with resulting truncated transcripts and no functional proteins.

### Culture of gastruloids

mES cells and gastruloid cultures were conducted as described in ref.^31^. mES cells were maintained in a humidified incubator (5% CO2, 37°C), and passaged every 3 days into 2i LIF DMEM medium composed of DMEM supplemented with 10% mES certified FBS, 100ng ml^−1^ of mouse LIF, 3μM of GSK-3 inhibitor (CHIR99021) and 1μM of MEK1/2 inhibitor (PD0325901). For gastruloid differentiation, 300 mES cells were resuspended in 40μl of prewarmed N2B27 medium (50% DMEM/F12 and 50% Neurobasal supplemented with 0.5x N2 and 0.5x B27), and seeded into each well of a low-attachment, rounded-bottom 96-well plate. For Chiron-treated gastruloid, after 48h, 150µl of N2B27 medium supplemented with 3μM of GSK-3 inhibitor was added to each well. For Activin A-treated gastruloid, 100ng ml^−1^ of recombinant Activin A was used instead of the GSK-3 inhibitor. Medium was then replaced every 24h. For experiments involving Matrigel, N2B27 medium was replaced at 96h with N2B27 containing 10% Matrigel. Collection of gastruloids for each timepoint was performed indiscriminately. Brightfield live imaging was performed using IncuCyte S3 (Sartorius). For quantification of the ectopic posterior structures, four independent batches were quantified and as having or not more than 2 structures per specimen (total gastruloids n = 76 wt versus 83 *Hox^−/−^*). The collective means for all batches were ca. 20% (wt) *versus* 60% (*Hox*-less). p-value was determined by unpaired two-sided t-test (**P < 0.01). Chiron and Activin A gastruloid size quantification was performed using ImageJ software and the spine length of each gastruloid was quantified by manual measurement.

### 2D endoderm differentiation

Endoderm differentiation *in vitro* followed the protocol from^72^. Around 50’000 cells of control and *Hox^−/−^* mES cells were seeded in a 6-wells plate in differentiation medium (DM) composed of DMEM/F12 with 0.2% FBS medium and 1x B27. After 24h, medium was replaced with DM supplemented with 100ng ml^−1^ of recombinant Activin A and 3μM of GSK-3 inhibitor. The same treatment was repeated at 72h, before harvesting cells at 120h.

### Whole mount *in situ* hybridization (WISH)

For WISH, 120h gastruloid were collected and processed as previously reported in ref^31^. Briefly, they were fixed overnight in 4% PFA at 4°C and stored in methanol at −20°C until ready for processing. Each sample was rehydrated and prepared with Proteinase K (EuroBio) at 2.5μg ml^−1^ for 2min. After post-fixation, gastruloid were prehybridized in a blocking solution at 68°C for 4h, before incubation overnight with specific digoxigenin-labeled RNA probes of *Cdx2* and *Meox1* (Supplementary Table 5), at a final concentration of 200ng ml^−1^. The next day, samples were washed and incubated with an anti-DIG antibody coupled to alkaline phosphatase (Roche, 1:3’000). Staining was performed with BM-Purple (Roche). Images of gastruloids were captured with an Olympus DP74 camera mounted on an Olympus MVX10 microscope using the Olympus cellSens Standard 2.1 software. For RNA FISH, gastruloids in Matrigel were collected and processed following the HCR method (Molecular Instruments). Samples were collected in HBSS and fixed 1h in 4% PFA at RT and sequentially dehydrated in methanol for storage at −20°C. Samples were rehydrated, treated with Proteinase K and postfixed as in the WISH procedure, then incubated overnight at 37°C in a hybridization solution containing a multiplexed mixture of *Pcdh8*, *Mesp2*, *Uncx* and *Tbx18* probes (Molecular Instruments) at 4nM each. The next day, gastruloids were washed and incubated overnight at RT in the amplification buffer containing a mixture of the corresponding hairpins for each probe at a concentration of 60nM. Samples were washed, mounted and imaged using a confocal microscope (Leica SP8).

### Immunofluorescence

Gastruloids cultured in Matrigel were collected at 120h in HBSS, washed and fixed 1h in 4% PFA at RT, then stored at 4°C until processed. Samples were washed and permeabilized in PBS containing 0.2% Triton X-100 (PBS-T) at RT followed by a 1h blocking step in PBS-T containing 10% FBS at 4°C. Gastruloids were incubated overnight at 4°C in the blocking solution containing the primary antibodies (anti-SOX2 1/100 (AF2018, R&D Systems); anti-T, 1/100 (ab20680, Abcam); anti-TBX6, 1/50 (AF4744, R&D Systems); anti-FOXC1, 1/100 (ab223850, Abcam); anti-CER1, 1/100 (MAB1986, R&D Systems); anti-SNAI1, 1/200 (C15D3, Cell Signaling). The next day, gastruloids were washed with PBS-T and incubated overnight at 4°C in the blocking solution containing the fluorescent secondary antibodies diluted at 1:500, Phalloidin (1:200, Alexa Fluo 488) and DAPI at 2µg ml^−1^. Samples were washed, mounted and imaged with confocal microscope (Leica SP8) using the same acquisition parameters for control and *Hox^−/−^*gastruloids. Counting of structure epithelialization with Phalloidin staining was performed using ImageJ software. For each gastruloid, visible Phalloidin condensations throughout the full z-stacks were counted manually.

### RT-qPCR

Control and *Hox^−/−^* gastruloid were collected at 120h and pooled into an Eppendorf tube, washed and stored at −80°C until RNA extraction. RNeasy Plus Micro kit (Qiagen) with on-column DNase digestion was used following the manufacturer’s instructions. Reverse transcription was performed with 1µg of purified RNA using the SuperScript VILO kit (Invitrogen). Real-Time PCR (Biorad) was performed using the GoTaq 2x SYBR Mix (Promega) with primers for *Rps9* (fw-GACCAGGAGCTAAAGTTGATTGGA, rv-TCTTGGCCAGGGTAAACTTGA), *Hoxa2* (fw-CTCGGCCACAAAGAATCCCTG, rv-TGTCTCTCGGTCAAATCCAGC), *Sox17* (fw- AGCAAGATGCTAGGCAAGTCT, rv- GCCGGTACTTGTAGTTGGGG) and *Foxa2* (fw- TTTAAACCGCCATGCACTCG, rv- ACGGAAGAGTAGCCCTCG). mRNA levels were referenced to *Rps9* using the delta delta Ct method^73^.

### Next-generation sequencing analysis and figure

All NGS analyses were performed on a local installation of galaxy^74^. The calculations for single-cell RNA-seq were performed using the facilities of the Scientific IT and Application Support Center of EPFL. All command lines and scripts to regenerate figures are available at https://github.com/hrekaik/scriptsForRekaikEtAl2025. Genomic tracks were plotted using pyGenomeTracks v3.7^75,76^ and modified with Illustrator 2024. All boxplots, bar plots, heatmaps, venn diagrams and figures relative to scRNA-seq, multiome or ChIPmentation were plotted with R (www.r-project.org), scCustomize^77^ was used to generate expression projection on UMAP.

### RNA-seq

The procedures for both RNA extraction and library preparation were conducted as described in^31^. Briefly, mES cells and endoderm differentiated mES cells were collected in RLT plus buffer (Qiagen) and stored at −80°C until RNA extraction. RNeasy Plus Micro kit (Qiagen) with on-column DNase digestion was used for RNA extraction. RNA-seq library preparation with Poly-A selection was performed with 1μg of purified RNA using the TruSeq Stranded mRNA kit from Illumina. Library quality was assessed with a fragment analyzer before sequencing on a AVITI sequencer as paired-end, 150bp reads. For data analysis using a local galaxy server (see above) adapter sequences were trimmed from reads using cutadapt^78^ v1.16 (-a ‘GATCGGAAGAGCACACGTCTGAACTCCAGTCAC’ -A ‘GATCGGAAGAGCGTCGTGTAGGGAAAGAGTGTAGATCTCGGTGGTCGCCGTATC ATT’). Trimmed reads were aligned on mm10 with STAR v2.7.7a^79^ with ENCODE options using a custom gtf based on Ensembl v102^80^. Only uniquely mapped reads (tag NH:i:1) were kept using bamFilter v2.4.1^81^ and Cufflinks^82^ v2.2.1 was run with default parameters to get FPKM values. DESeq2 package was used to produce the differential gene expression^83^. The genes were considered as significant when the adjusted p-value was below 0.05 and the absolute log2 fold change above 1.5.

### ATAC-seq

ATAC-seq dataset from control and *Hox^−/−^* mES cells were produced as described in^84^. Cells were dissociated with Accutase for 5min at 37°C, washed and collected in cold resuspension buffer (RSB). 50k cells were resuspended in lysis buffer and incubated for 3min on ice. then immediately washed with RSB containing 0.01% Tween-20 and centrifuged. The pellet was gently resuspended in 50μL Tagmentation Mix (Nextera) and incubated at 37°C in a thermomixer at 1000rpm for 30min. DNA was extracted using the MinElute PCR Purification Kit (Qiagen), then fragments were amplified with Nextera Index Primers (Nextera) using the KAPA HiFi HotStart ReadyMix (Roche) for 12 cycles. After PCR the reactions were cleaned first with the MinElute PCR Purification Kit and then with AMPure XP beads (Beckman Coulter). Library quality was assessed with a fragment analyzer before sequencing on a AVITI sequencer as paired-end, 150bp reads. Adapter sequences and bad quality bases were removed with Cutadapt^78^ v4.8 with options -a CTGTCTCTTATACACATCTCCGAGCCCACGAGAC -A CTGTCTCTTATACACATCTGACGCTGCCGACGA -q 30 -m 15. Reads were mapped with bowtie^85^ v2.5.0 with parameters --very-sensitive on mm10. Alignments with a mapping quality below 30, discordant pairs, and reads mapping to the mitochondria, were discarded with bamtools v2.5.2 [https://github.com/pezmaster31/bamtools]. PCR duplicates were removed with picard v2.18.2 [http://broadinstitute.github.io/picard/index.html] before the BAM to BED conversion with bedtools^86^ v2.30.0. Coverage and peak calling were computed by MACS2^87^ v2.2.9.1 with options --format BED --gsize 1870000000 --call-summits --keep-dup all --bdg --nomodel --extsize 200 --shift −100. Coverages were normalized to million reads in peaks (summits +-500bp) using bigwigAverage from deeptools^76^ v3.5.4.

### Single-cell RNA-seq

scRNA-seq assay was performed as described in^88^. Control and *Hox^−/−^* gastruloid at 72h, 96h 120h and 144h were collected, dissociated and processed following the manufacturer’s instructions of the 3’ CellPlex Kit (10X Genomics). Cells were then pooled and subjected to single-cell RNA-seq using the 10x Genomics platform (V3.1 chemistry). cDNA preparations were performed according to 10X Genomics recommendations, amplified for 10–12 cycles and sequenced on a Novaseq 6000 with the cbot2 chemistry. For each condition, except at 144h and wt at 72h, two replicates were performed. Fastqs of one wt 96h sample were retrieved from GEO (GSM7890954 and GSM7890936)^89^ and processed with datasets generated in this study. For data processing, a Galaxy pipeline was used to preprocess the 10x CellPlex libraries: Mapping and demultiplexing of cell barcode were performed using RNA STARSolo v2.7.10b against the mouse reference genome mm10 and a modified gtf file based on Ensembl 102^90^ with the following parameters --soloBarcodeReadLength 1 --soloCBstart 1 --soloCBlen 16 --soloUMIstart 17 --soloUMIlen 12 --soloStrand Forward --soloFeatures Gene --soloUMIdedup 1MM_CR --soloUMIfiltering - --quantMode TranscriptomeSAM GeneCounts --outSAMattributes NH HI AS nM GX GN CB UB --outSAMtype BAM SortedByCoordinate --soloCellFilter None --soloOutFormatFeaturesGeneField3 ‘GeneExpression’ --outSAMunmapped None --outSAMmapqUnique 60 --limitOutSJoneRead 1000 --limitOutSJcollapsed 1000000 --limitSjdbInsertNsj 1000000. Then, the matrices were filtered with DropletUtils^91,92^ v1.10 (Method EmptyDrops, lower-bound threshold = 100, FDR threshold = 0.01). Samples demultiplexing was performed using CITE-seq-Count (https://zenodo.org/badge/latestdoi/99617772) v1.4.4 --cell_bacode_first_base 1 --cell_barcode_last_base 16 --umi_first_base 17 --umi_last_base 28 --bc_collapsing_dist 1 --umi_collapsing_dist 2 --expected_cells 24000 --whitelist ‘3M-february-2018.txt’ --max-error 2. Subsequently the cell barcodes were translated to match the one from the gene expression. Each gene–cell count matrix was then analyzed using Seurat v5.1.0^93^. A Seurat object was generated by filtering out barcodes with less than 200 identified gene and genes identified in less than three cells. Demultiplexing from the translated CMO matrices was then performed: any cell barcode with less than 5 CMO UMI was excluded as well as any cell barcode absent from the Seurat object. Sample attribution was performed with demuxmix^94^ using the total number of gene expression UMI per cell. Additional filtering was applied to remove low-quality cells and doublets. For each dataset, the mean RNA count was used and cells with either less than 40% of the mean, or more than 2.5-fold of the mean were removed. In addition, cells with more than 8, or less than 0.05% mitochondrial counts were removed (Supplementary Table 6). Then, all individual datasets were merged into a single Seurat object. The combined dataset was then normalized and the 2000 most variable features were identified and data scaled and regressed by cell cycle score and percentage of mitochondrial reads. Principal component analysis (PCA) was computed using variable genes falling within the 5th and 80th percentile of expression and by excluding genes included in the regions of the deletions of the four *Hox* clusters. UMAP and k-nearest neighbours were then computed using 30 principal components. Clusters were computed at a resolution of 1.5 in order to identify the NMP population. Clusters with similar identities (using known gene markers) were then manually grouped together. For the differential gene expression analysis, the function FindMarkers from Seurat was used and the thresholds were set to adjusted p-value below 0.01 and absolute log2 fold-change above 1.5. For RNA velocity, velocyto v0.17.17^95^ was used to pre-processes datasets and generate loom files. scVelo v1.10.2^96^ was used to estimate and project RNA velocity and compute the pseudotime. The velocity was computed once for the whole dataset in order to sort cells by pseudotime in the heatmap, and a second time only on the *Hox^−/−^* cells to project the pseudotime on the UMAP. In order to compare the dataset to a single-cell atlas, matrices from E6.5 to E9.5 were downloaded from https://tome.gs.washington.edu^32^. The E9.5 sample was downsampled to get only 20’000 cells. All mouse samples were merged with the Seurat object of gastruloids and the gene labels were unified. The cells were split by study^32,97–99^, normalized and 2500 variable features were identified. Genes included in the regions of the deletions of the four *Hox* clusters were excluded. Data was scaled and PCA was run with 50 components. Finally, all cells from published studies^32,97–99^ and ours were integrated with the IntegrateLayers function from Seurat with CCAIntegration, recursively. UMAP was computed using the 30 first component of the integrated cca.

### Multiome analysis

Nuclei isolation for the single-cell Multiome ATAC and gene expression procedure was performed following the manufacture’s instruction of the Chromium Nuclei Isolation Kit (10X Genomics). Gastruloid at 96h and 120h were collected, pooled and dissociated as described in the scRNA-seq procedure. Dissociated cells were resuspended in cold lysis buffer and incubated for 10min on ice, transferred to the nuclei isolation column and centrifuged at 16’000 rcf for 20sec at 4°C to remove cellular debris. The Flowthrough was centrifuged to retrieve the nuclei pellet, then washed with cold Debris Removal Buffer and two rounds of centrifugation, and wash with cold Wash Buffer. Purified nuclei were resuspended in cold Resuspension Buffer, counted and their nuclear membrane assessed. Samples were immediately processed according to 10X Genomics recommendations of the v1.0 Multiome ATAC + Gene Expression kit. The 10X ATAC and GEX libraries were sequenced on a Novaseq 6000 instrument. scRNA data was analysed as described above to generate a Seurat object except that the valid barcode files were 737K-arc-v1.txt. For scATAC data processing, first, the I2 fastq was reverse complemented with seqtk seq v1.3.3. Then cell barcodes were added to fastq reads using Sinto barcode v0.10.1. After adapters removal with cutadapt v4.9, mapping was performed with BWA-MEM v0.7.18 against the mouse reference genome mm10. ATAC fragments with the associated cell barcode were retrieved with Sinto fragments v0.9.0. The barcodes were translated to match the one from the gene expression. These translated fragment files were sorted and bgzipped to be used by Signac. On the other hand, the fragment file was duplicated and attributed to the + and - strand in order to use both extremities in the peak calling. This duplicated fragment file was used to call peaks with MACS2^87^ v2.2.9.1 with options --format BED --gsize 1870000000 --call-summits --keep-dup all --bdg --nomodel --extsize 200 --shift −100. The peaks from wt and from *Hox^−/−^* samples were concatenated, sorted and merged with bedtools v2.31.1 in order to create a single comprehensive peak list. Downstream analysis was done in R with Signac v1.14.0^100^. Individual chromatin assays were created using the ATAC fragment object and the comprehensive list of peaks, then added to the RNA Seurat object. wt and *Hox^−/−^* Seurat objects were then merged. The combined Seurat object was filtered to remove low quality cells using ATAC counts (below 1800 or above 100’000), RNA counts (below 1’000 and above 25’000), nucleosome signal (above 2) and TSS enrichment (below 1) values. RNA layer was normalized with SCTransform regressing the percentage of mitochondrial genes and the cell cycle. PCA was also performed on selected genes using the average expression, excluding *Hox* genes. For the ATAC layer, all features with more than 5 counts were considered as top features and data normalized using the RunTFID function. The singular value decomposition (SVD) was computed after exclusion of genomic intervals spanning the deleted *Hox* regions. The joint neighbor graph using RNA and ATAC assays was performed with the FindMultiModalNeighbors function, with the 50 first PC for SCT (RNA) and the 2^nd^ to 40^th^ for lsi (ATAC). The UMAP was computed using this ‘weighted.nn’ and clustering was computed using the grah ‘wknn’ with a resolution of 0.5. For motif analysis, chromVAR v1.26.0^101^ was used with the JASPAR 2022 and JASPAR 2024 motif collection^102^, the motifs from ref^103^, the ISMARA motif collection^104^ and the HOMER motifs. The pseudotime was computed as for the single-cell RNA-seq in order to sort the cells in the heatmap.

#### ChIPmentation

ChIPmentation experiments were performed as described in^31^. Briefly, collected gastruloid were pooled in a 15ml falcon tube, washed with PBS and resuspended in 1ml PBS containing 1% formaldehyde for fixation during 10 min at RT. The crosslink reaction was stopped by adding a glycine solution to a final concentration of 0.125M. Fixed gastruloids were pelleted and stored at −80°C until further use. Samples were resuspended in a sonication buffer (Tris HCl pH=8.0 50mM; EDTA 10mM; SDS 0.25% and protease inhibitors) and sonicated in a Covaris S220 device for 14 min (duty cycle 2%, peak incident power 105W) to obtain an average chromatin fragment size of 300-500 bp. A dilution buffer (HEPES pH=7.3 20mM; EDTA 1mM; NP40 0.1%; NaCl 150mM and protease inhibitors) was added to the sonicated chromatin and incubated with the antibody-bead complex (Pierce Protein A/G Magnetic Beads, Thermo Scientific with 5µg anti-PBX1 (4342S, Cell Signaling)) overnight at 4°C. Sequential washes were then performed twice with RIPA buffer (Tris HCl pH=8.0 10mM; EDTA 1mM; Sodium Deoxycholate 0.1% TritonX-100 1%; NaCl 140mM and protease inhibitors), RIPA High salt buffer (Tris HCl pH=8.0 10mM; EDTA 1mM; Sodium Deoxycholate 0.1% TritonX-100 1%; NaCl 500mM and protease inhibitors), LiCl buffer (Tris HCl pH=8.0 10mM; EDTA 1mM; LiCl 250mM; Sodium Deoxycholate 0.5%; NP40 0.5% and protease inhibitors) and Tris HCl buffer (pH=8.0 10mM and protease inhibitors). DNA fragments bound to the antibody-bead complex were tegmented using the Nextera Tegmentation kit. Beads were resuspended in the tegmentation buffer and incubated at 37° for 2min with 1µl of the Tn5 transposase. Fragment were then eluted and purified as described previously, and amplified using Nextera primers. Final DNA libraries were purified and size selected using AMPure XP magnetic beads (Beckman Coulter), and a fragment analysis was performed before sequencing on a DNBSEQ-G40 sequencer as paired-end, 100bp reads. For data analysis, adapter sequences and bad quality bases were trimmed from reads using cutadapt ^78^ version 1.16 (-a ‘CTGTCTCTTATACACATCTCCGAGCCCACGAGAC’ -a ‘CTGTCTCTTATACACATCTGACGCTGCCGACGA’ --quality-cutoff=30). Trimmed reads were mapped on mm10 genome using bowtie2 version 2.3.4.1^105^. Only alignments with proper pairs and mapping quality above 30 were kept using samtools 1.8. Peaks and coverage were obtained with macs2 version 2.1.1.20160309 (--format BAMPE --gsize 1870000000 --call-summits --bdg). Motif analysis of PBX1 ChIPmentation peaks was performed with HOMER version 4.10. ChIPmentation quantifications and heatmaps were performed with deeptools version 3.5.6^76^.

## ACKNOWLEDGEMENTS

We thank all members of the Duboule laboratories, as well as Angela Nieto and Alfonso Martinez-Arias, for comments and discussions. The calculations were performed using the facilities of the Scientific IT and Application Support Center of EPFL. This work was supported in part using the resources and services of the Gene Expression Research Core Facility (GECF) at the School of Life Sciences of EPFL.

## AUTHORS CONTRIBUTIONS

Conceptualization: HR, DD

Methodology: HR, LLD, AM

Investigation: HR, LLD, YR, CD, AMC, JG

Funding acquisition: DD

Project administration and supervision: DD

Writing the original drafts: DD, HR

Review writing & editing: AM, LLD

## FUNDING

This work was supported by funds from the Ecole Polytechnique Fédérale (EPFL) Lausanne, the University of Geneva, the Collège de France (Paris), the Swiss National Research Fund (No. 310030_196868 and CRSII5_189956).

## COMPETING INTERESTS

The authors declare that they have no competing interests.

## DATA AVAILABILITY

All raw and processed datasets are available in the Gene Expression Omnibus (GEO) repository under accession number GSE288886.

## CODE AVAILABILITY

All scripts necessary to reproduce figures from raw data are available at: https://github.com/hrekaik/scriptsForRekaikEtAl2025

## LEGENDS TO EXTENDED DATA FIGURES

**Extended Data Figure 1.**
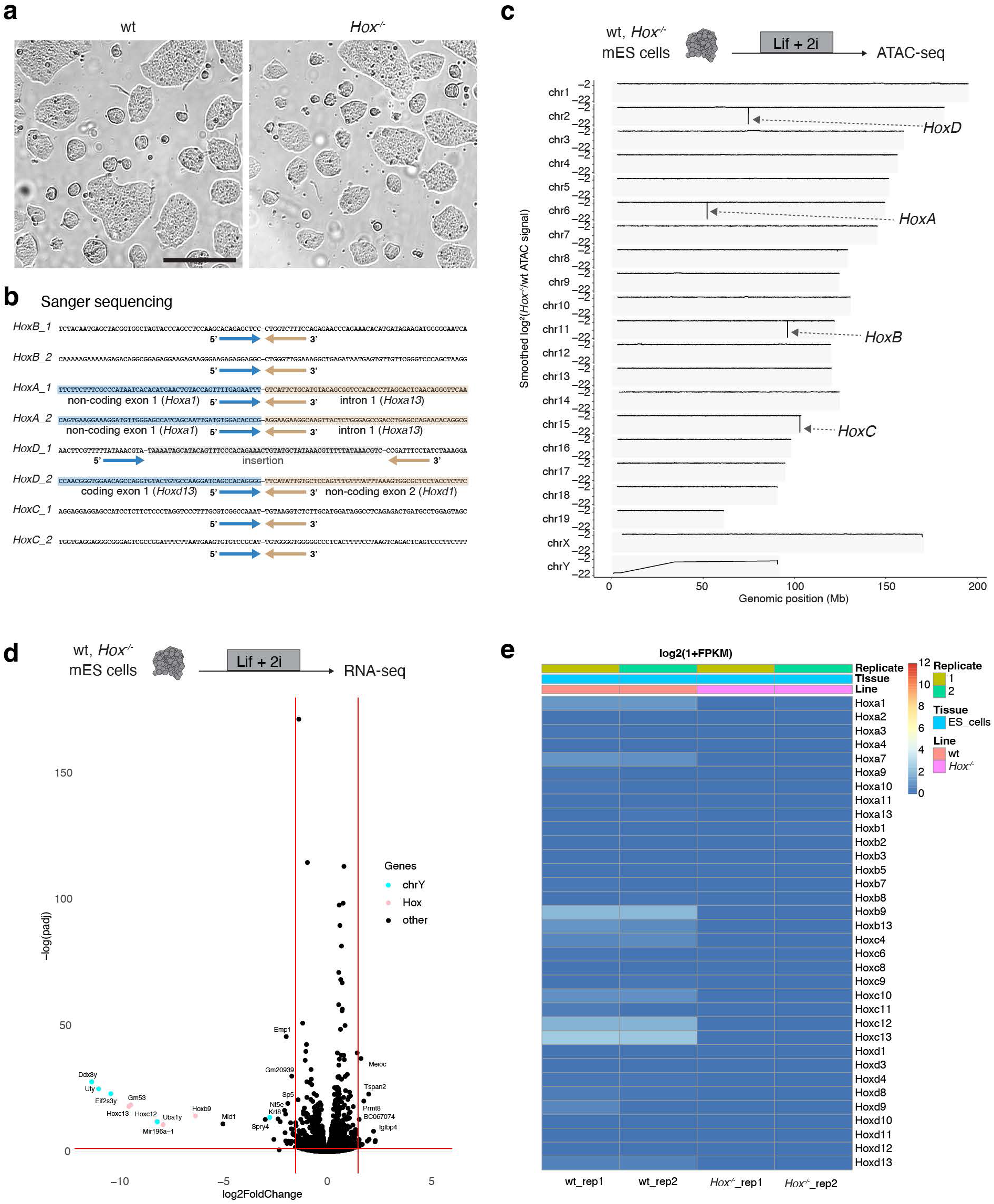
Characterization of both wt and *Hox^−/−^* mES cells by ATAC-seq and RNA-seq. **a;** Representative brightfield images of wt and *Hox^−/−^* ES cells. Cells were maintained in Lif + 2i medium to prevent cell differentiation. Scale bar: 100 µm. **b;** Schematic representation of sanger sequencing results showing the deletions produced in each allele of the four *Hox* clusters. Blue and ochre arrows indicate the 5’ and 3’ regions around each breakpoint, respectively. Sequences are highlighted when insertions are present or when they fall within gene annotations. Both mutant alleles of the *HoxA* cluster contain the second exon of *Hoxa13*, while the second allele of the *HoxD* cluster contains a partial region of the first *Hoxd13* exon. In all of these alleles, no functional protein is produced. **c;** Genome-wide analysis using ATAC-seq data from wt and *Hox^−/−^* cells, assessing potential deletions present in the mutant compared to wt, for each chromosome (n = 2). **d**; Volcano plot of the RNA-seq dataset to show genes differentially expressed between wt and *Hox^−/−^* mES cells. Each point represents one gene and vertical and horizontal red lines represent the threshold of a 1.5 log2 fold change variation and the adjusted p-value < 0.05. In addition to genes located within the *Hox* clusters (in pink), several genes located within the Y chromosome were also lost (in cyan, n = 2). **e;** *Hox* gene expression in wt and *Hox^−/−^* ES cells. While no expression of *Hox* genes was detected in the mutant, only weak levels of expression for a few *Hox* genes were observed in control ES cells.

**Extended Data Figure 2.**
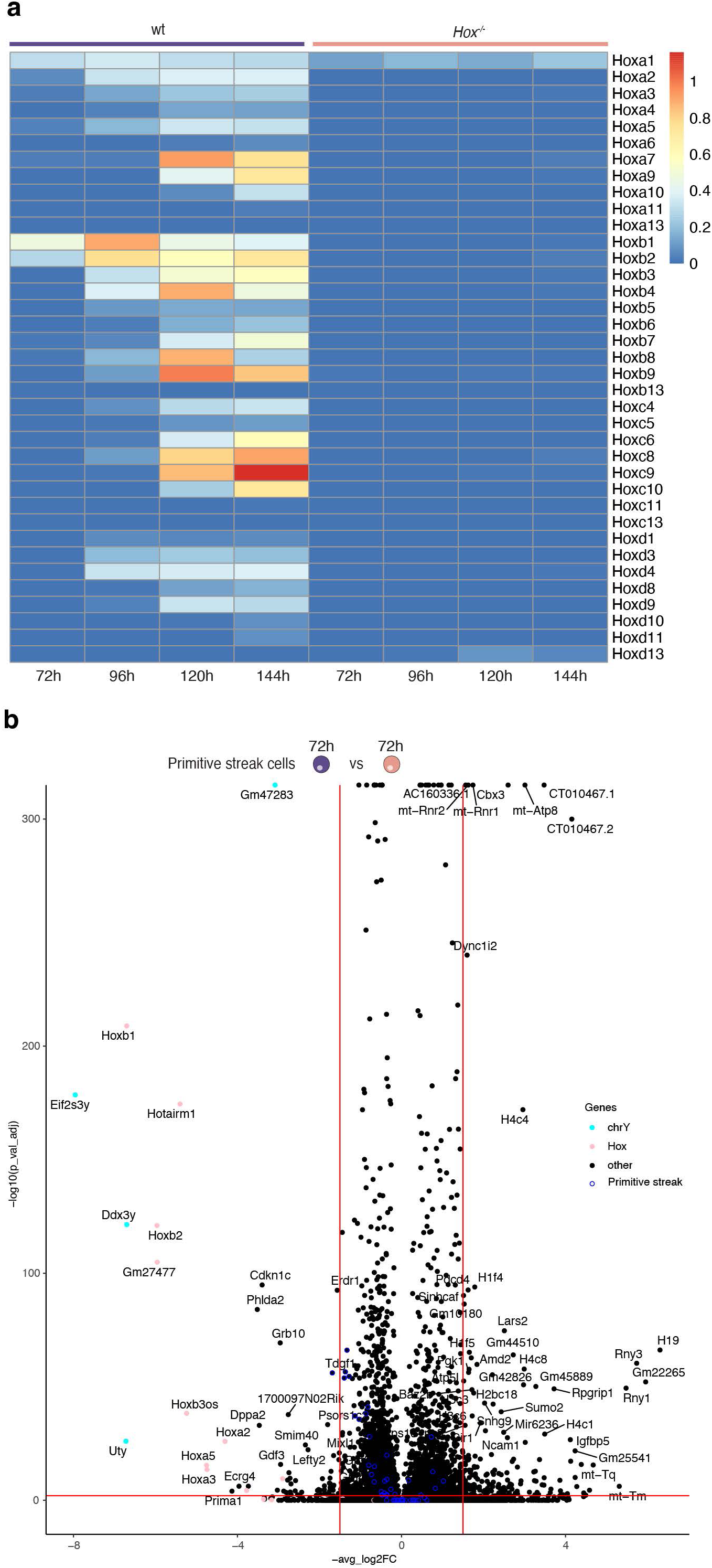
scRNA-seq analysis of gastruloids. **a;** Heatmap of *Hox* genes expression in control (wt) and *Hox^−/−^* mutant gastruloids at 72h, 96h, 120h and 144h. For each gene, the average expression in all cells at the same timepoint was calculated. Expression in wt follows temporal colinearity. **b;** Volcano plot of differentially expressed genes between wt and *Hox^−/−^* specimens in primitive streak cells of 72h gastruloids. Each point represents one gene and vertical and horizontal red lines represent the threshold of a 1.5 log2 fold change variation and the adjusted p-value < 0.05, respectively. The top 50 markers of the primitive streak are highlighted in blue, with no significant change observed. Few *Hox* genes (in ochre) were expressed in this cellular population.

**Extended Data Figure 3.**
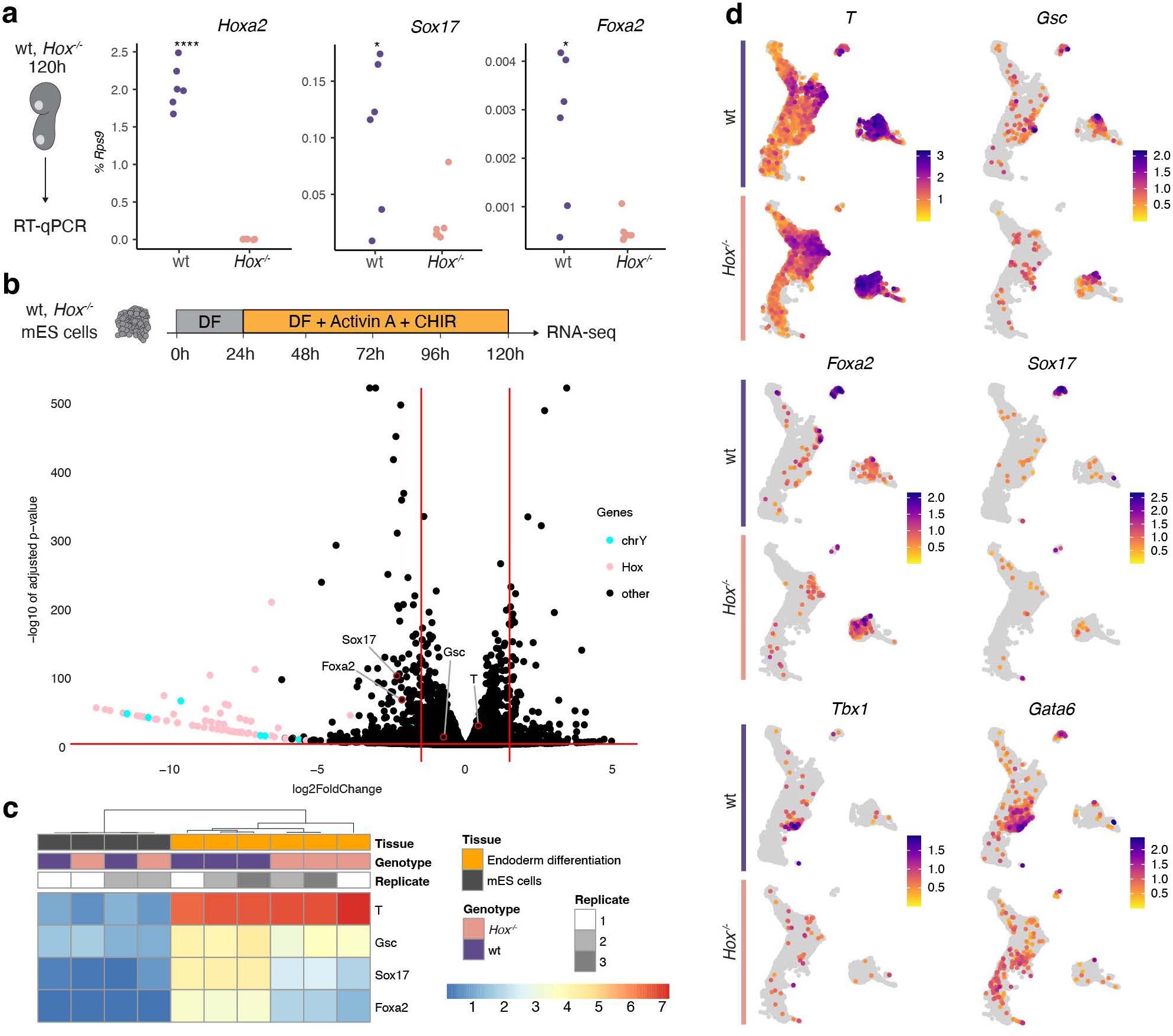
Endoderm is undetectable from either *Hox^−/−^* gastruloids, or in an *in vitro* endoderm differentiation protocole. **a;** RT-qPCR of conol (wt) and *Hox^−/−^*gastruloids at 120h showing very low level of endodermal markers in the mutant condition. Expression of genes is calculated relative to *Rps9* (n = 6). *P* values were determined by unpaired two-sided *t*-test (\**P* < 0.05 and \*\*\*\**P* < 0.0001). **b;** *In vitro* endodermal differentiation of wt and *Hox^−/−^* mES cells using an Activin A + CHIR protocol (top). Cells were analyzed by RNA-seq (n = 3). Below, the corresponding Volcano plot shows those genes differentially expressed between the two conditions (bottom). Each point represents one gene and vertical and horizontal red lines represent the threshold of a 1.5 log2 fold change variation and the adjusted p-value < 0.05. Red dots highlight marker genes for either anterior primitive streak (*T* and *Gsc*), or endodermal differentiation (*Sox17* and *Foxa2*). wt and *Hox^−/−^*differentiated cells showed similar level of anterior primitive streak markers, whereas expression of endodermal markers was significatively decreased in mutant cells. **c;** Heatmap of anterior primitive streak and definitive endoderm genes in mES cells RNA-seq shown in Extended Data Fig. 1 and the endoderm differentiation RNA-seq shown in **b**. In the differentiation protocol towards endoderm, cells shows a significant decrease in the expression of endodermal markers in the mutant condition. **d;** scRNA-seq UMAP of wt and *Hox^−/−^*gastruloids highlighting markers of either anterior primitive streak (*T* and *Gsc*), endodermal differentiation (*Sox17* and *Foxa2*), or <u>cardio</u>pharyngeal mesoderm (*Tbx1* and *Gata6*). Expression of anterior primitive streak markers is scored in mutant gastruloids. In contrast, neither endoderm, nor <u>cardio</u>pharyngeal mesoderm markers were detected in *Hox^−/−^* gastruloids. UMAP are the same as in Figure 2.

**Extended Data Figure 4.**
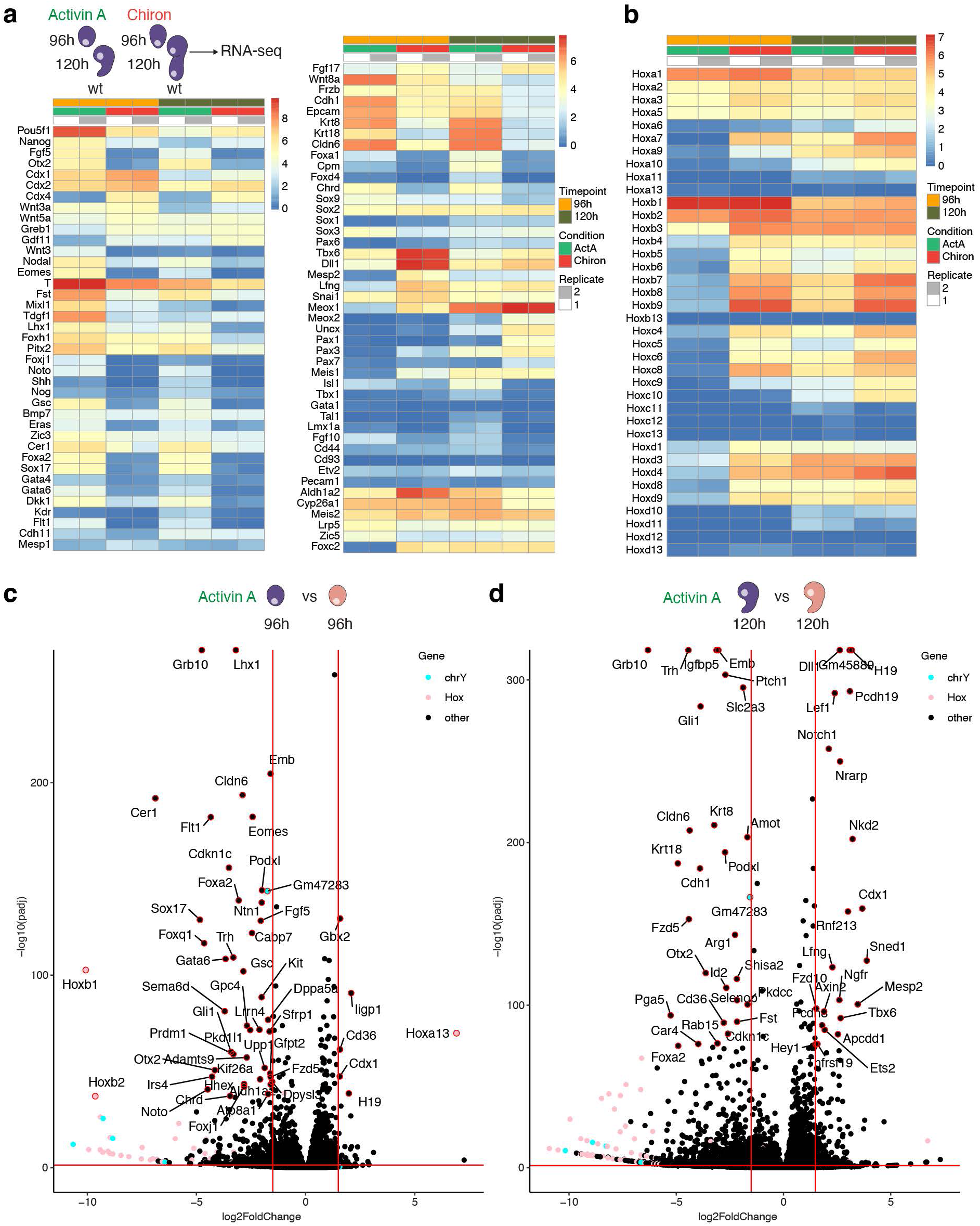
Transcriptome analysis of gastruloids treated with either Activin A or Chiron. **a;** Schematic design for RNA-seq analysis of Activin A- or Chiron-treated gastruloids at 96h and 120h, produced from wt mES cells. This comparison assesses differences between the two *in vitro* systems. Heatmap of some markers selected from ref.^35^ confirms that the Activin A protocol generates more anterior tissues (e.g., enhanced endoderm/cardiopharyngeal markers), when compared to the Chiron protocol (n = 2 replicates per condition). **b;** Heatmap showing the dynamic expression of *Hox* genes in Activin A- versus Chiron-treated gastruloids using wt cells at 96h and 120h. **c-d;** Volcano plot of differentially expressed genes between wt and *Hox^−/−^* Activin A-treated gastruloids at 96h (c) and 120h (d). Vertical and horizontal red lines indicate the threshold of 1.5 log2 fold change and adjusted p-value < 0.05, respectively. Pink dots highlight *Hox* genes, and cyan dots indicate genes located on the Y chromosome (n = 2 replicates).

**Extended Data Figure 5.**
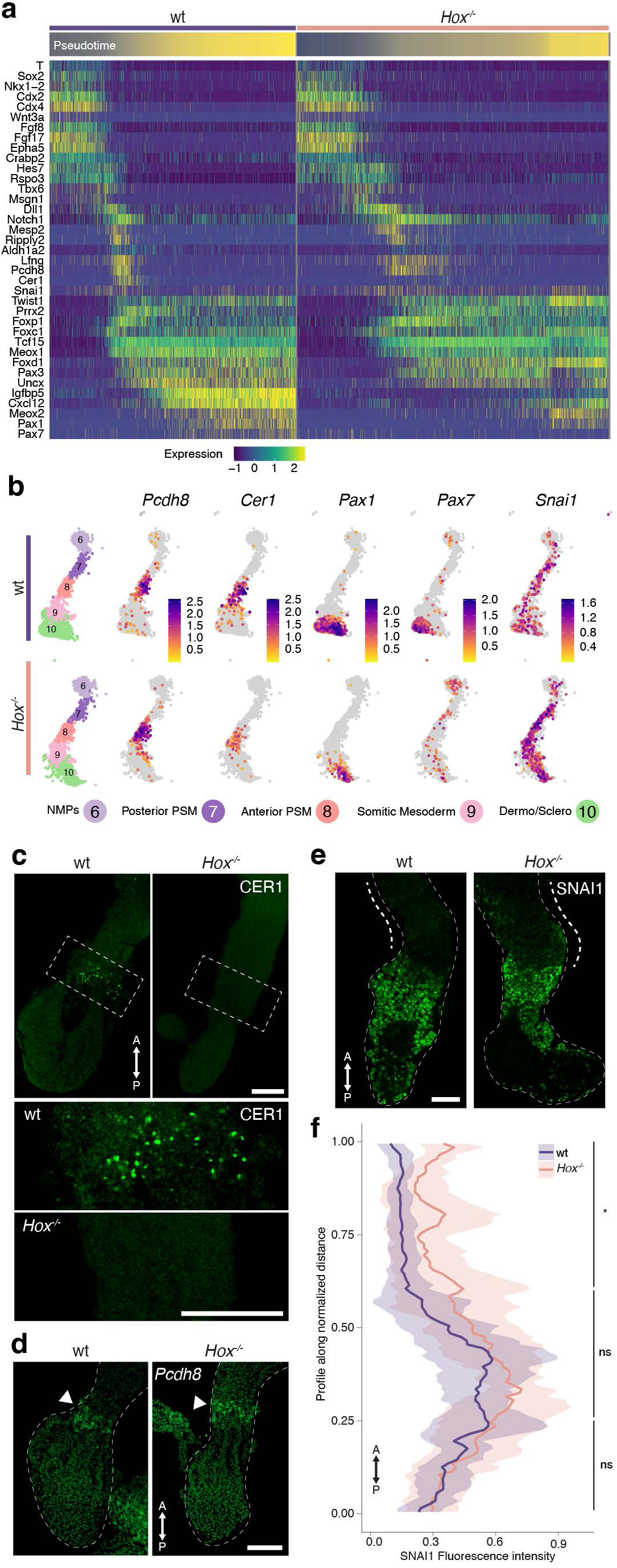
Expression of markers of the NMPs to somite differentiation trajectory. **a;** Expression heatmap of selected genes using the scRNA-seq data extracted from cells participating to the NMP to somitic trajectory at 120h and 144h. The expression of each gene in individual cells (n = 2666 and 3373 cells in wt and *Hox^−/−^* condition, respectively), as displayed along the x axis and ordered by their pseudotime along the differentiation trajectory, showed similarity in both NMPs and somitic markers between control and *Hox^−/−^* cells. However, *Cer1* expression as well as mature dermomyotome marker (*Pax7*) were severely decreased in mutant cells. **b;** UMAP projection emphasizing the expression of some markers (top) along the NMPs to somite differentiation trajectory at 120h and 144h, with the corresponding clusters (left, bottom). Unlike *Pax1*, *Pax7* expression was lost in mutant mature somites. Expression of *Pcdh8*, specific to the anterior part of the forming somite, was maintained in mutant cells, suggesting that the down-regulation of *Cer1* was likely not due to the loss of these cells in mutant gastruloids. In contrast, *Snai1* expression was upregulated in the somitic tissue. **c-g**; Spatial validation of RNA expression changes. **c;** Immunofluorescence of CER1 in 120h control and *Hox^−/−^*, matrigel-exposed gastruloids. Staining shows the location of the excreted CER1 protein at the PSM to SM transition in control gastruloids, while absent from the mutant counterpart. Scale bar: 200 µm. **d;** HCR for *Pcdh8* in control and *Hox^−/−^*gastruloids at 120h. Scale bar: 100 µm. **e;** Immunofluorescence of SNAI1 in 120h control and *Hox^−/−^* gastruloids. Scale bar: 100 µm. Staining shows the weak yet significant maintenance of the staining within somitic tissue of the mutant (dashed line). **g;** Quantification of the distribution of SNAI1 staining along the anterior-posterior axis of 120h gastruloids. The profile (fluorescence intensity along normalized distance) is shown for control and *Hox^−/−^* specimens as a line with a ribbon (mean ± SD, n = 8). A significance increase was observed in the somitic region of mutant gastruloids. p-value is determined by Wilcoxon rank-sum test on individual sections (*P < 0. 05).

**Extended Data Figure 6.**
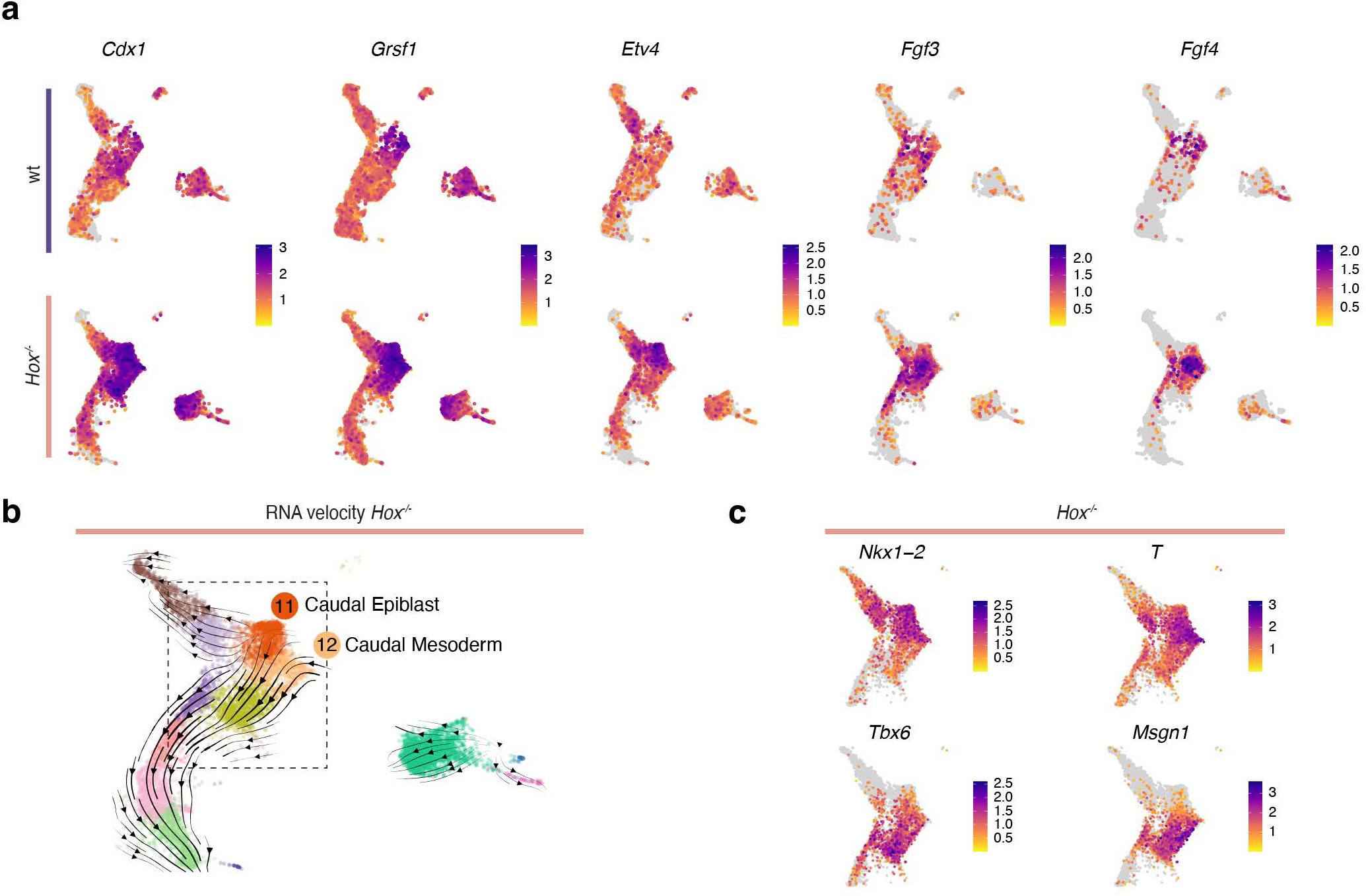
Comparative scRNA-seq of caudal epiblast and caudal mesoderm cells. **a;** scRNA-seq UMAP of control (wt) and *Hox^−/−^* gastruloids showing the expression of selected genes expressed either in caudal epiblast, or in caudal mesoderm clusters, with a specific increase in intensity observed in *Hox^−/−^*specimens. UMAPs are as in Figure 2. **b;** scRNA-seq UMAP of *Hox^−/−^* gastruloids with projected RNA velocity. Streamlines represent the velocity vector field that estimates the probabilities of cell-to-cell transitions. Colors represent the clusters as in Figure 2d. A transition between caudal epiblast and caudal mesoderm cells was observed. **c;** UMAP of *Hox^−/−^* gastruloids displaying the expression of markers for the differentiation process illustrated in **b** (dashed box). Expression of progenitor markers (i.e., *Nkx1-2* and *T*) in the caudal epiblast cells was progressively replaced by the expression of mesodermal markers (i.e. *Tbx6* and *Msgn1*), in caudal mesoderm cells.

**Extended Data Figure 7.**
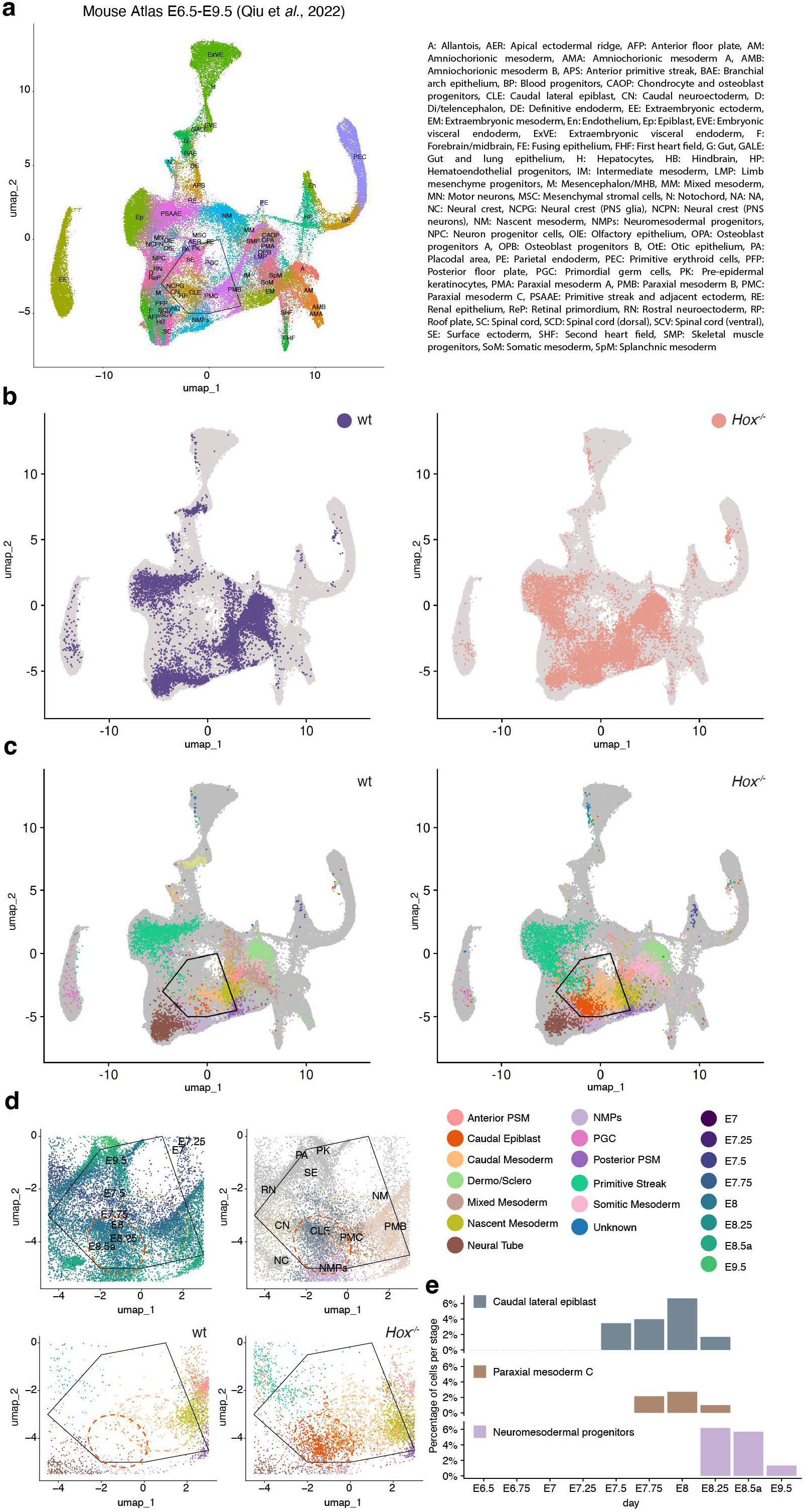
Projections of control and mutant gastruloid cells on UMAP of mouse embryos. **a;** scRNA-seq of mouse embryos between E6.5 and E9.5 on the integrated UMAP projection. Data were extracted from refs^97–99^ and reanalyzed. Colors represent cell type reported in ref^32^, with the legend on the right. **b, c;** UMAP integration of mouse embryos and gastruloids cells. Control (wt, left) and *Hox^−/−^* (right) gastruloids cells are highlighted within the integrated UMAP. In **c,** the colors represent cluster identities of gastruloids cells, as shown in Figure 2d. Caudal epiblast and caudal mesoderm cells were enriched in mutant specimen and are highlighted within a black polygon. **d;** Magnified view of the embryo UMAP showing embryonic cells on top, with colors reflecting either the embryonic stages (upper left with legend on the right), or the atlas cell type as in **a** (upper right, legend in panel **a**, right). UMAPs at the bottom show gastruloids cells (wt, left and *Hox^−/−^*, right), with colors representing their identities (as in **c**; legend on the upper right). The red and orange circles highlight the caudal epiblast and caudal mesoderm populations, respectively. Mutant caudal epiblast cells were comparable to the CLE (caudal lateral epiblast) cells of the embryo, whereas mutant caudal mesoderm cells resembled the embryonic paraxial mesoderm type C cells. These two embryonic populations were normally detected at ca. embryonic stage E7.5 to E8.25. **e;** Proportion of embryonic CLE cells, paraxial mesoderm type C cells and NMPs cells at each embryonic stage. CLE cells are detected in the embryo almost one day before the appearance of NMPs cells, when the former start to disappear.

**Extended Data Figure 8.**
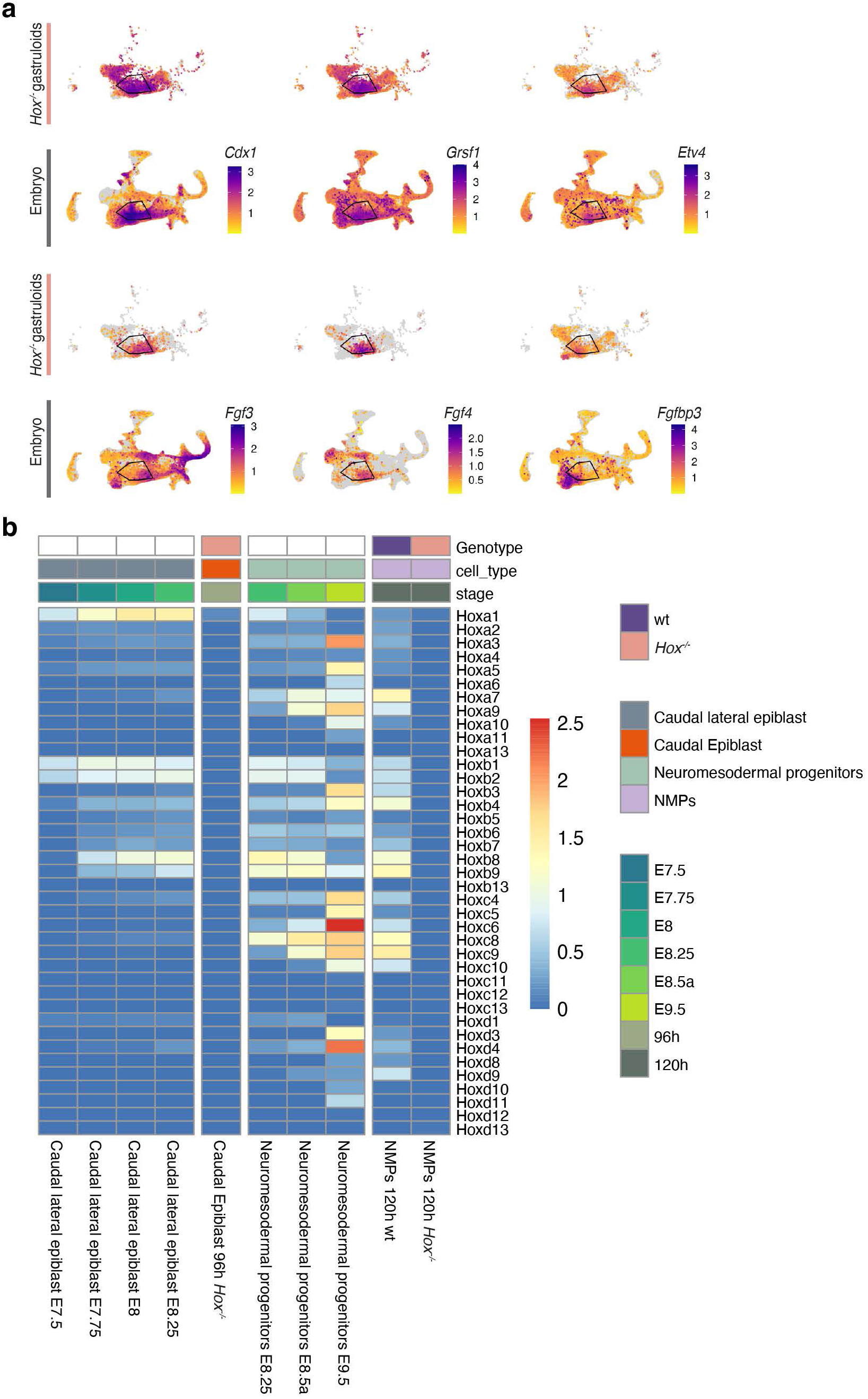
Comparative expression of selected genes in *Hox^−/−^* gastruloids and murine embryos. **a;** scRNA-seq integrated UMAPs of *Hox^−/−^* gastruloids (top) and mouse embryos (bottom). Expression of selected markers, which were identified as overexpressed in mutant gastruloids, within the caudal epiblast and caudal mesoderm clusters (black polygon). Equivalent expressions were observed within the embryonic caudal lateral epiblast (CLE). **b;** Heatmap of *Hox* gene expression in gastruloids and mouse embryos in caudal epiblast (left) and NMP (center) populations. Only few *Hox* genes were expressed in early mouse embryonic CLE cells (i.e., at E7.5), comparable to the expression in caudal epiblast population of *Hox^−/−^*gastruloid. Subsequent stages of CLE cells showed more *Hox* genes expressed, in particular *Hoxa* and *Hoxb* genes. In contrast, comparable patterns and levels (with a few exceptions) of *Hox* gene expression were observed between either embryonic or control gastruloid NMP cells.

**Extended Data Figure 9.**
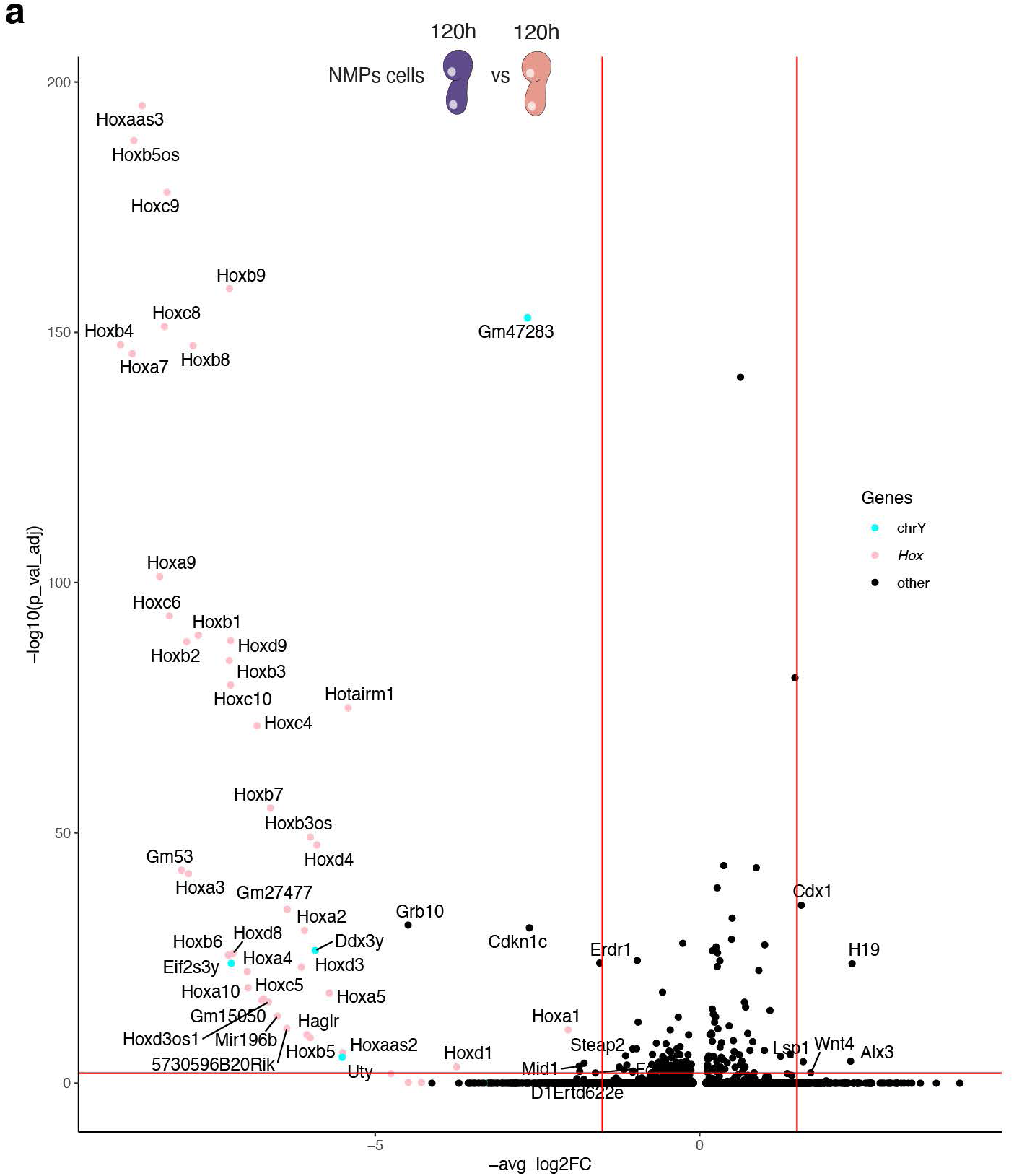
Gene expression in NMP cells. **a,** Volcano plot of differentially expressed genes between control (wt) and *Hox^−/−^* NMP cells from 120h gastruloids. Each point represents one gene, vertical and horizontal red lines represent the threshold of a 1.5 log2 fold change variation and the adjusted p-value < 0.05, respectively. Besides *Hox* genes themselves and some LncRNA genes located within *Hox* clusters (pink dots), only very few other genes were affected by the absence of any HOX proteins, even at 120h, i.e., days after the initiation of *Hox* gene expression in NMP cells.

**Extended Data Figure 10.**
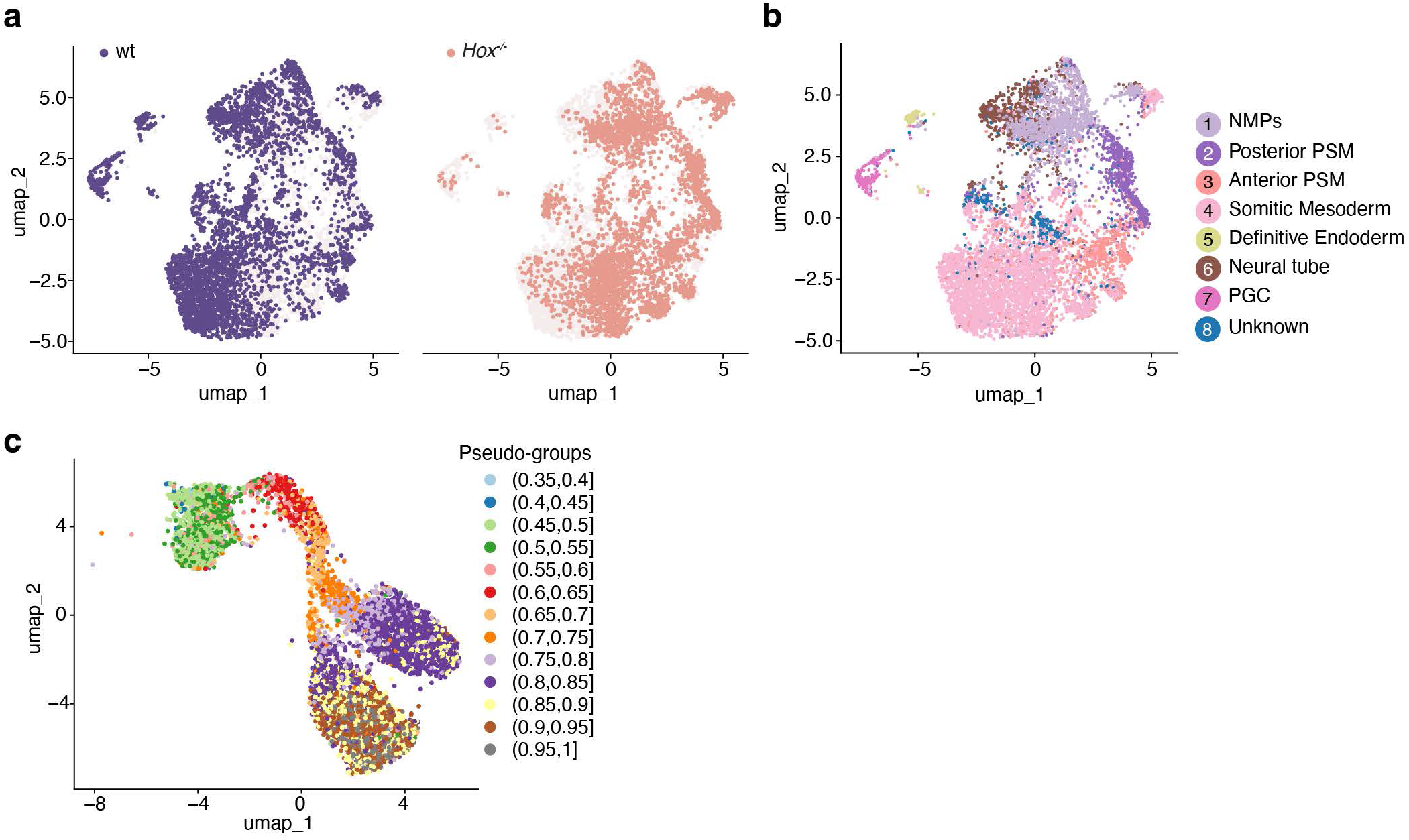
Multiome analysis of 120h gastruloids. **a;** UMAP of multiome data at 120h using RNA projection, with cells colored by genotype (wt, purple; *Hox^−/−^,* orange), showing the distribution of cells. **b;** RNA UMAP projection showing cells colored by unsupervised clusters, with cluster identities indicated on the right. **c;** UMAP of multiome data with cells colored according to their pseudotime-based pseudo-groups.

## SUPPLEMENTARY INFORMATION

**Supplementary Figure 1:**
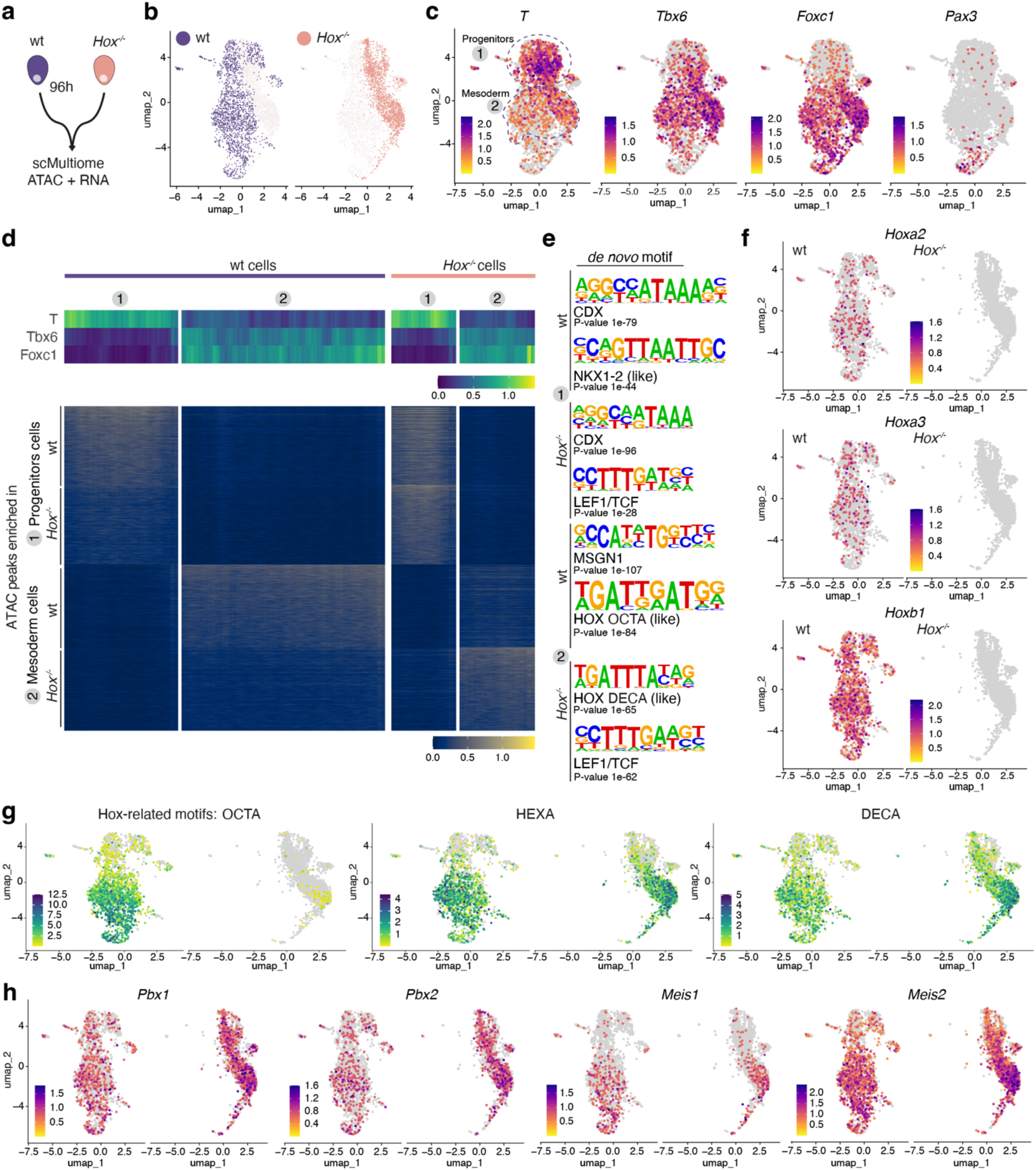
Multiomics comparative analysis of 96h control and *Hox*-less mutant gastruloids. **a;** 96h control and *Hox^−/−^* gastruloids were used to generate a single cell multiome dataset, combining both ATAC and RNA analysis. **b;** Projection of the integrated multiome UMAP, with cells colored by their genotype (wt left, purple; *Hox^−/−^* cells right, orange). **c;** UMAPs showing the expression of selected genes specific either to progenitor cells (upper dashed circle labelled 1), or to differentiating mesoderm (lower dashed circle labelled 2). **d;** Heatmap of control and *Hox^−/−^* cells clustered by their identity (x-axis) and showing RNA expression of the same markers as in **c** (top), as well as chromatin accessibility using ATAC peaks (bottom). The top 500 peaks enriched either in progenitor or in mesodermal cells, in both control and *Hox^−/−^* cells, were quantified. In the progenitor population (1), the lack of any HOX function led to decreased accessibility. In contrast, in differentiating mesodermal cells, specific ATAC peaks were observed for each genotype. **e;** Homer motif analysis of the top 500 peaks extracted from each cell population quantified in panel **d**. Displayed are the top 2 *de novo* binding motifs for each cell population, with the most probable matches to known motifs (below). In progenitors, LEF1/TCF motif was enriched in *Hox^−/−^* cells. In mesodermal cells, HOX OCTA motif was enriched in control cells, whereas HOX DECA motif was enriched in mutant counterparts. **f,** Multiome UMAP showing similar expression (RNAs) of some anterior *Hox* genes in both progenitor and mesodermal cells. **g;** Multiome UMAP showing chromVAR motif activity of selected motifs present in ATAC peaks (see Figure 5). The accessibility pattern of the HOX OCTA motif was much enriched in control *versus Hox^−/−^* mesodermal cells, as observed in **e**, whereas the HOX DECA motif appeared slightly enriched in a majority of mutant cells. **h;** Multiome UMAP showing *Pbx1*, *Pbx2*, *Meis1* and *Meis2* RNA expression in both progenitor and mesodermal cells. Expression is observed throughout all cell types in both control and mutant gastruloid cells, except for *Meis1* RNAs, which were restricted to differentiating mesoderm cells.

**Supplementary Movie 1**

Elongation of wt and *Hox^−/−^* gastruloids in Matrigel between 96h and 120h. Somitic-like condensations are produced in the control specimen (left, black arrowheads), while they are not detected in the mutant (open black arrowheads). In contrast, several outgrowths are observed in the posterior region of *Hox^−/−^*gastruloids (white arrowheads). Scale bar = 400 µm.

**Supplementary Table 1.** Differential expression of genes between control and *Hox^−/−^* cells within the NMPs to somite differentiation trajectory at 120h.

This table is provided as a separate Supplementary Excel file.

**Supplementary Table 2:**
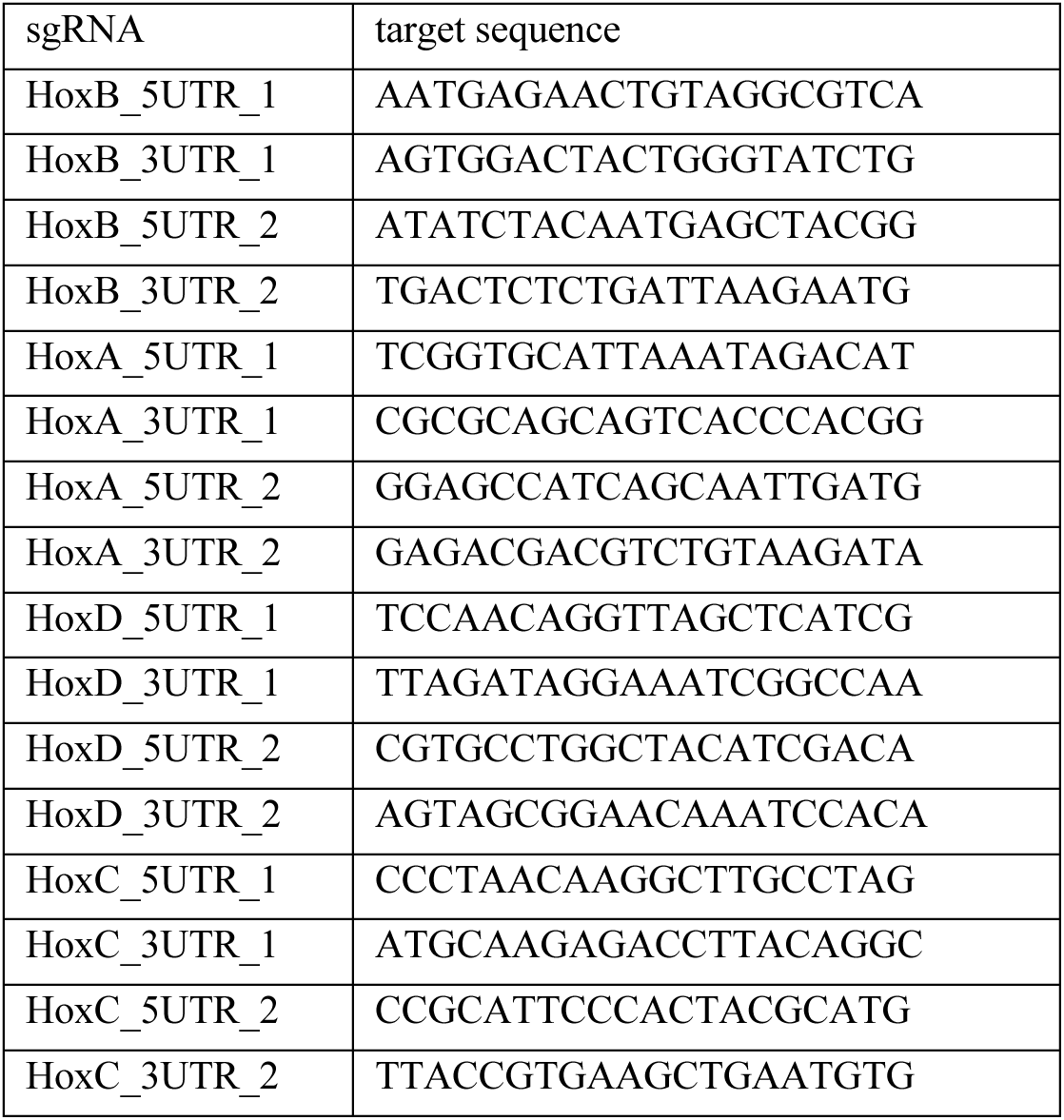
List of sgRNA sequences used for *Hox* cluster deletions.

**Supplementary Table 3:**
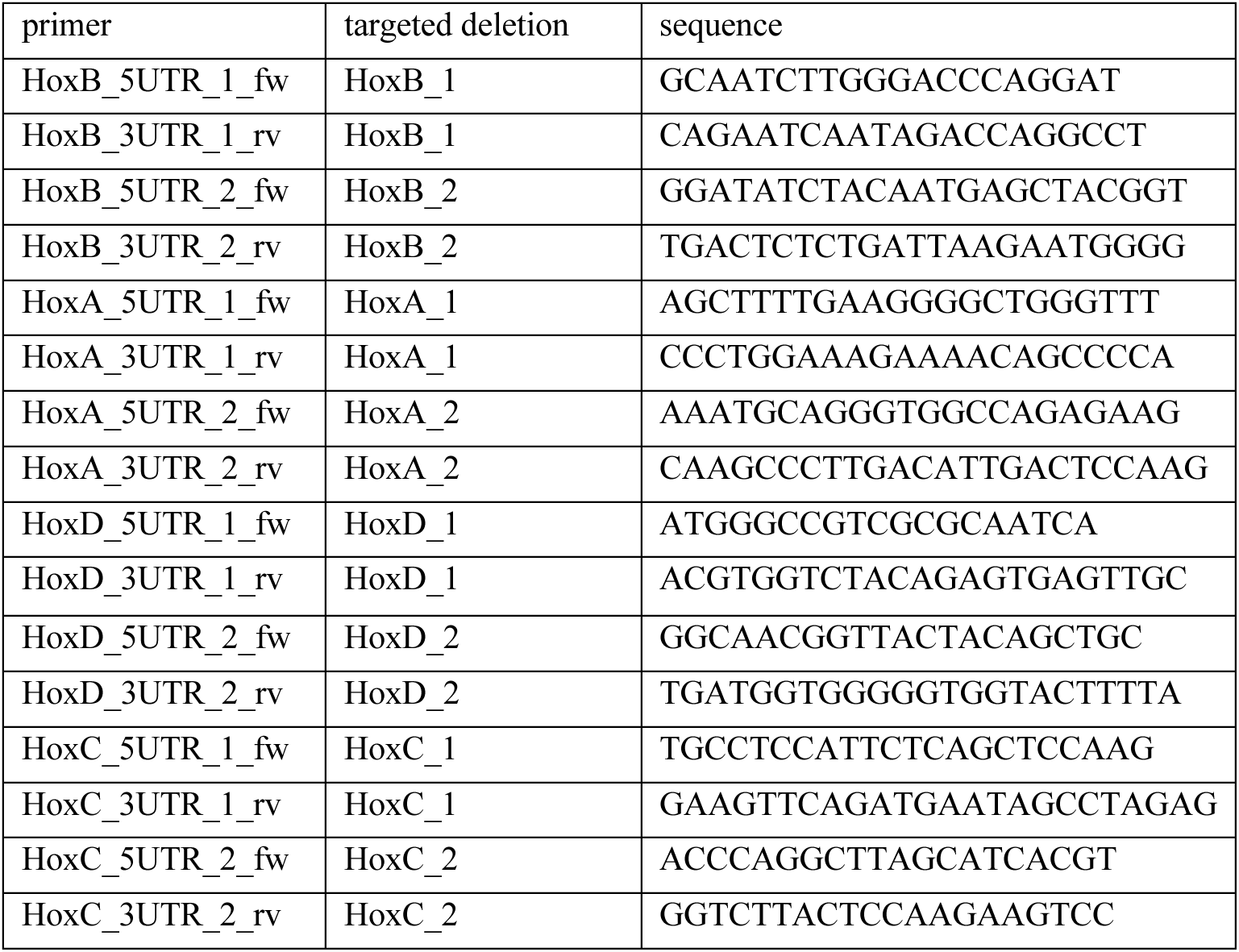
List of primers used for genotyping.

**Supplementary Table 4:**
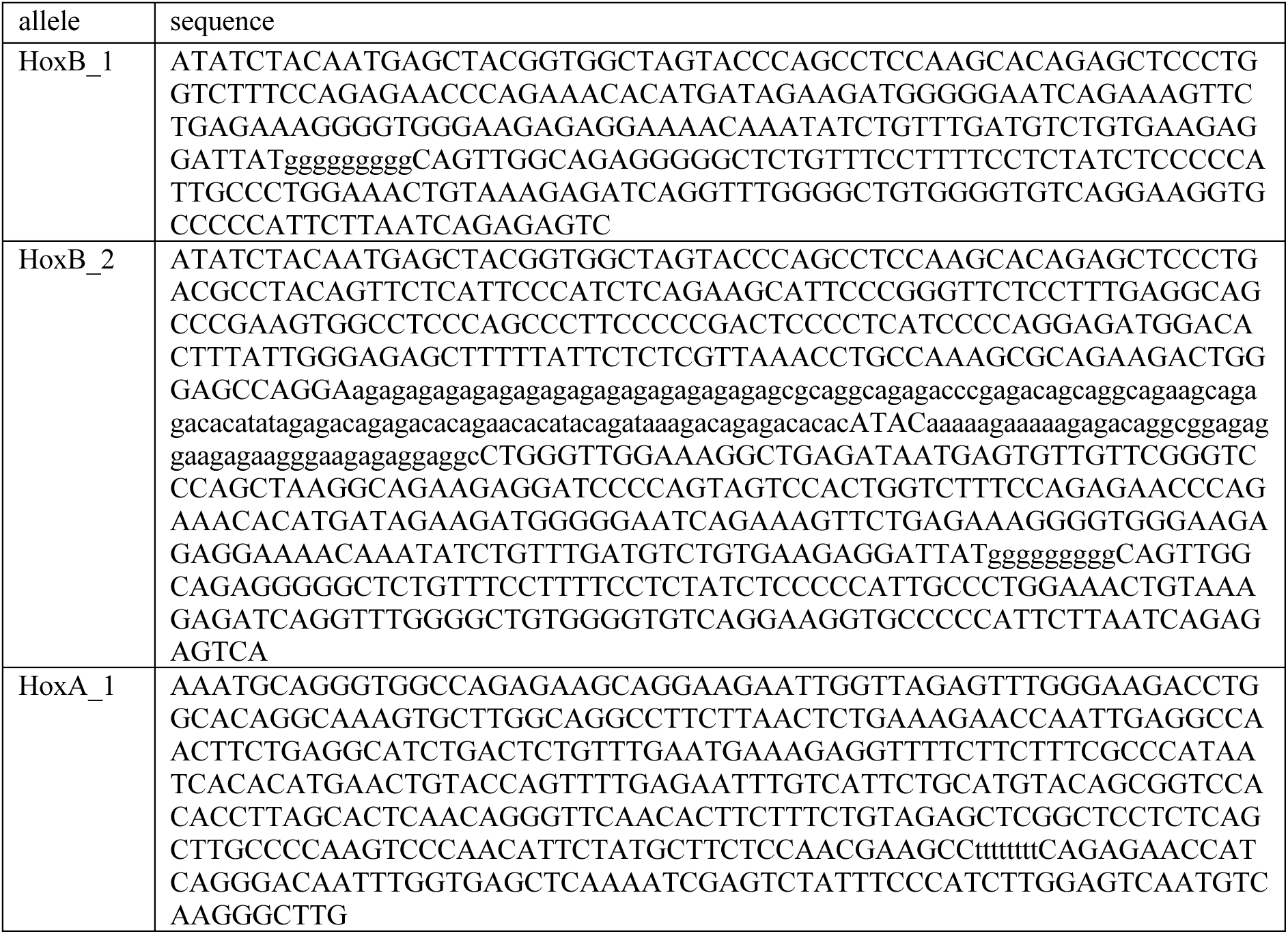
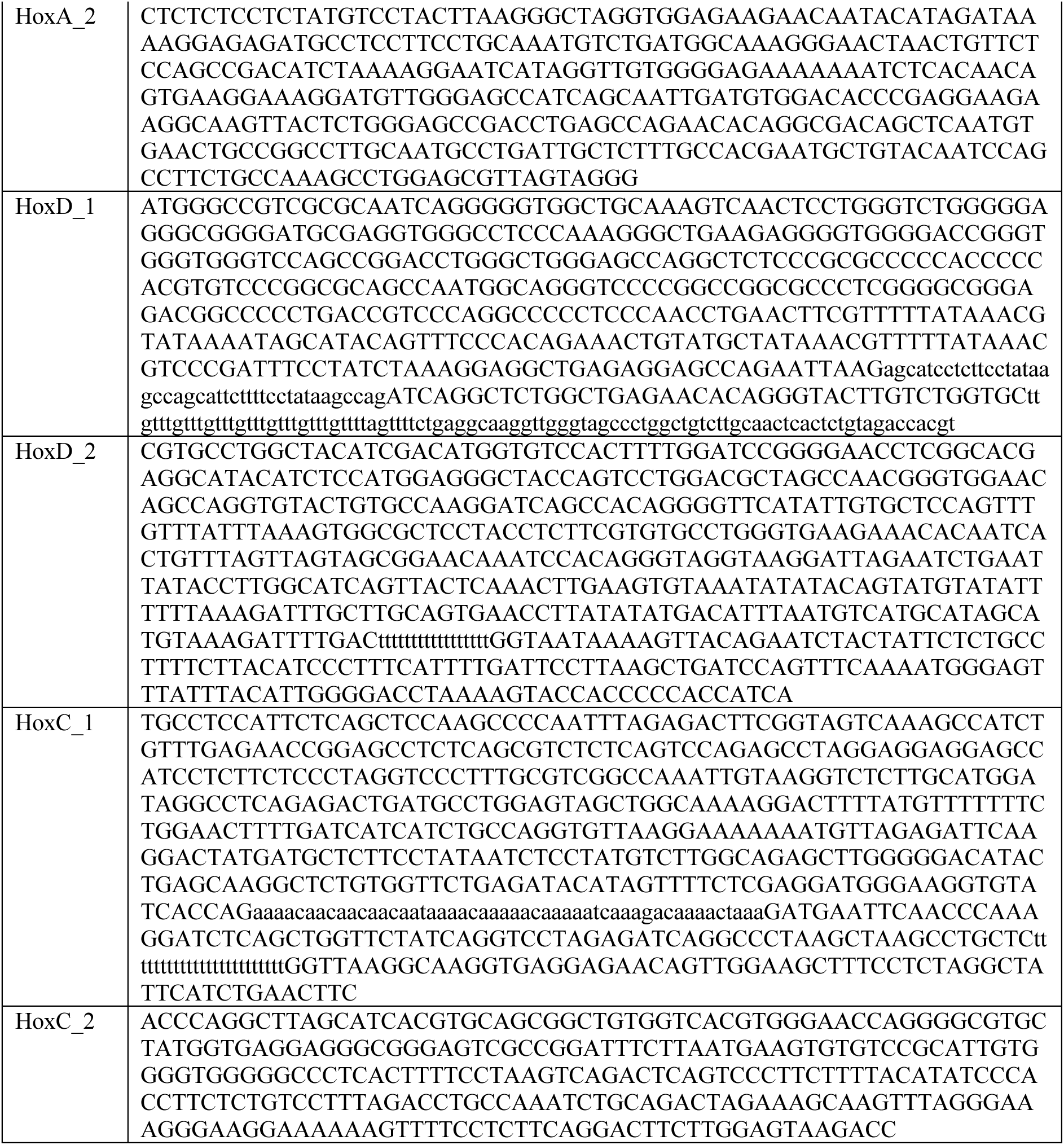
Sequence of *Hox* cluster deletions in the *Hox^−/−^* mES cell line.

**Supplementary Table 5:**
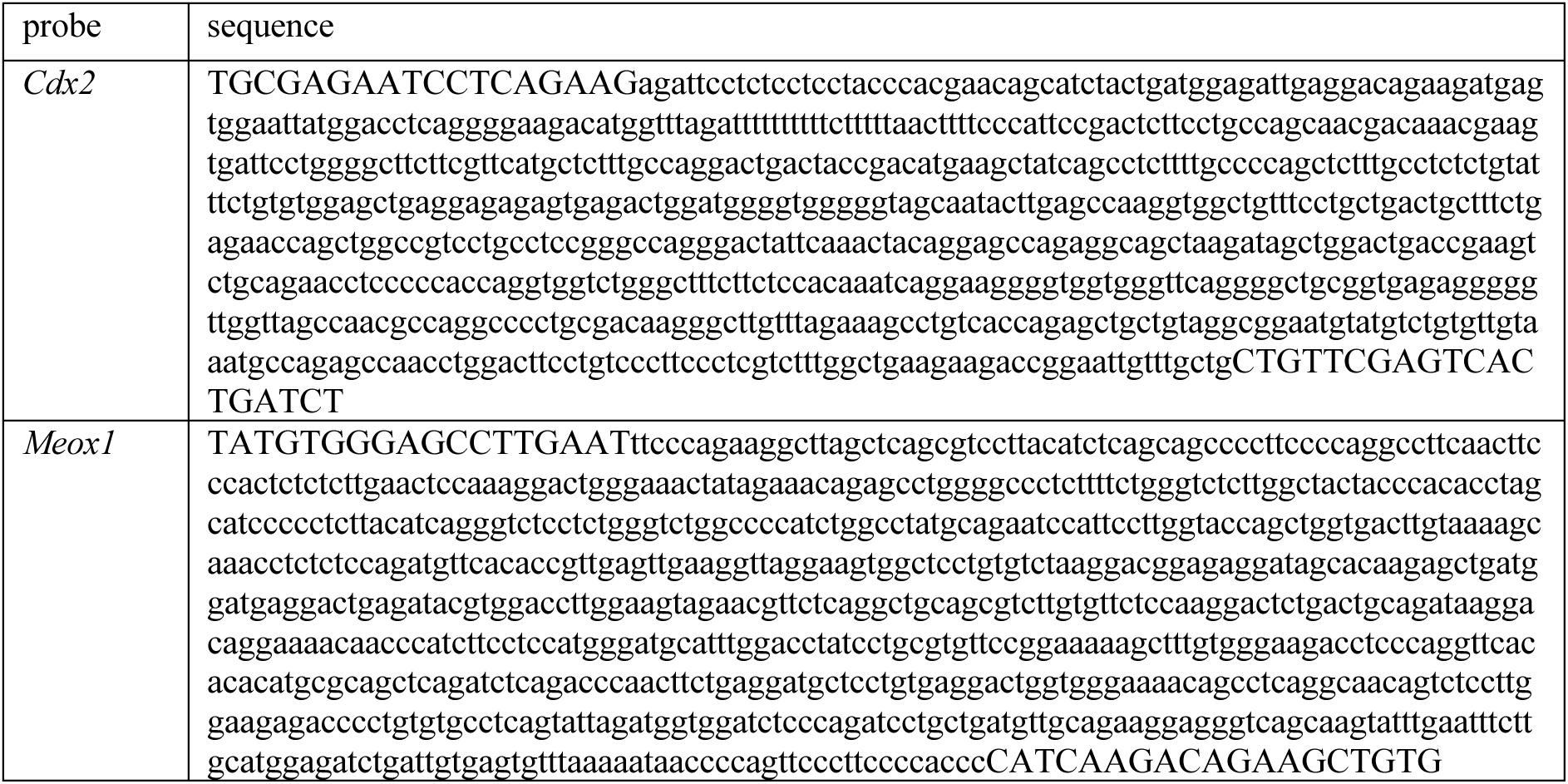
Sequence of WISH RNA probes.

**Supplementary Table 6:**
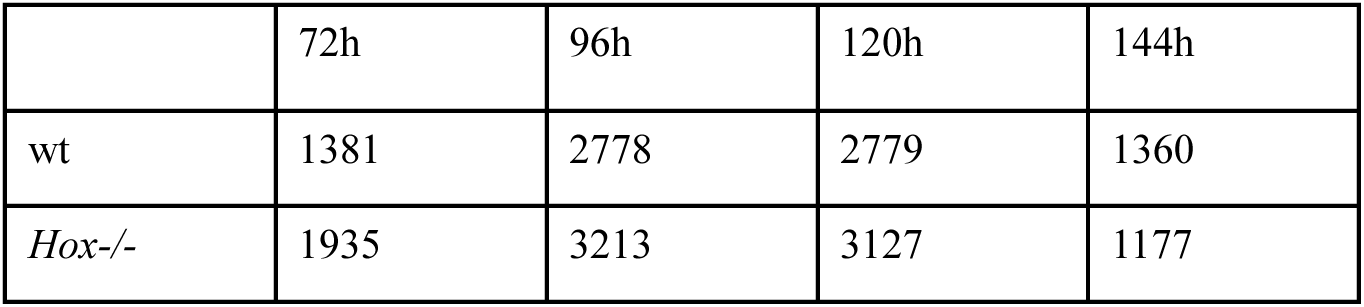
Number of scRNA-seq cells retained after filtering for each condition and time point:

|  | 72h | 96h | 120h | 144h |
| --- | --- | --- | --- | --- |
| wt | 1381 | 2778 | 2779 | 1360 |
| *Hox*<sup>-/-</sup> | 1935 | 3213 | 3127 | 1177 |

## Notes

### Competing Interest Statement

The authors have declared no competing interest.

